# CYCLON is a nucleolar protein that regulates pan-cancer cell fitness through ribosome biogenesis

**DOI:** 10.64898/2026.05.13.724782

**Authors:** Andrea C. Garcia-Sandoval, Sébastien Durand, Nic Robertson, Sieme Hamaidia, Jan Mikolajczyk, Fleur Bourdelais, Emilie Montaut, Victoria Lopez, Tomasz Turowski, David Tollervey, Jean-Jacques Diaz, Olivier Destaing, Anouk Emadali

## Abstract

Tumor progression is driven by cancer cell fitness, defined as the capacity of malignant cells to maintain growth, adapt to stress and withstand therapy Cellular fitness is fundamentally governed by nucleolar processes, which act as central regulators by integrating RNA processing with ribosome biogenesis to support protein synthesis and stress adaptation. The nuclear protein CYCLON, containing a large intrinsically disordered region (IDR) could be implicated in mediating biomolecular condensates and regulatory plasticity, which are key elements of nucleolar biology. CYCLON also emerged as a candidate regulator of cancer cell fitness, as it is frequently overexpressed across tumor types. Inducible silencing and cell biology approaches have shown that CYCLON maintains nucleolar integrity, controls nucleoli size and number, nucleolin and Ki-67 distribution and prevents nucleolar stress. CYCLON contributes to ribosome biogenesis by binding ribosomal RNA (rRNA) and regulating 32S and 21S pre-rRNA processing, ultimately influencing ribosomal subunit production and global protein synthesis. Its depletion impairs proliferation and clonogenic capacity by prolonging both interphase and mitosis, leading to slowed cell cycle progression. The impact of CYCLON on cellular fitness has been consistently observed across cancer models, reinforcing its essential role in the regulation of nucleolar biology.

## Introduction

Cellular fitness refers to a cell’s ability to survive and proliferate by integrating its capacity to withstand environmental stress, acquire and utilize resources. This capacity is sustained by specific interplay between anabolic processes such as protein synthesis, DNA replication and associated checkpoints, as well as catabolic energy metabolism (1–3). Despite their central role in cellular physiology, the molecular determinants connecting these cellular activities remain poorly characterized.

CYCLON/CCDC86 is a nuclear protein that may participate in this interplay, but remains poorly characterized. A network of evidence suggests that CYCLON could be essential for development and cancer, as gene knock-out leads to embryonic lethality in mice (4), while depletion induces apoptosis in HeLa cells (5) and reduces cell survival in most cancer cell lines (DepMap database) (6). Moreover, CYCLON expression has been reported to be associated with tumor progression and immunotherapy resistance in cellular and mice models of B-Cell lymphoma and more specifically Diffuse Large B Cell Lymphoma where it is strongly associated with poor prognosis (7, 8). CYCLON depletion has also been recently shown to lead to mitotic defects in HeLa cells (5) and impair nasopharyngeal cancer invasiveness though EGFR and PI3K/AKT signaling (9). Under normal conditions, CYCLON is predominantly expressed in highly proliferative contexts such as testis and activated immune cells. It was shown to be induced in B-cell lymphocytes upon IL-3 stimulation (10) and to participate in Fas-dependent apoptosis in T-cell lymphocytes after TCR stimulation (4). CYCLON has also been described as a MYC target gene in both physiological (10, 11), and cancer-associated settings (5, 7). The involvement of CYCLON in development, immune regulation and oncogenesis underscore the necessity of a comprehensive functional analysis to define molecular mechanisms governing its essential functions.

CYCLON is a predominantly intrinsically disordered protein whose only clearly defined structural feature is a putative C-terminal coiled-coil RNA-binding domain, that shares homology with yeast Cgr-1, a protein implicated in nucleolar integrity and ribosome biogenesis (12). CYCLON has a reported nucleolar localization in lymphoma (13) and was shown to interact with several nucleolar-resident proteins including nucleophosmin/NPM1 (13), fibrillarin and Ki-67 (5) (IntAct and BIOGRID) as well as DNA and RNA (14–21).

The nucleolus is a nuclear domain primarily responsible for ribosome biogenesis. This process involves the transcription and processing of ribosomal RNA (rRNA) as well as the coordinated assembly of ribosomal proteins into functional ribosomal subunits (22). Consequently, nucleolar dynamics is central to cellular anabolism, supporting high levels of protein synthesis required for growth, proliferation and adaptation to changing physiological conditions (22). Beyond its canonical role in ribosome production, the nucleolus also serves as a regulatory hub for various cellular stress responses (23), cell cycle control (24), and heterochromatin organization (25). Nucleolus therefore represents a key determinant of cellular fitness through its ability to maintain or adapt vital cellular functions in response to intrinsic factors and external stresses (26).

CYCLON is expressed in cells with rapid proliferation, requiring elevated levels of protein synthesis and optimized metabolism, consistent with a role in promoting nucleolar activity. We hypothesized that CYCLON has an evolutionary conserved function in ribosome biogenesis and assessed this in human cancer cell lines. Functional investigation demonstrated that CYCLON is required to maintain nucleolar structure and regulates ribosome biogenesis. Notably, its depletion was sufficient to trigger the nucleolar stress response. CYCLON depletion results in reduced clonogenic potential and proliferation capacities by increasing the duration between cell divisions. These properties of CYCLON appear to represent a general mechanism promoting cancer cell fitness across multiple tumor types.

## Materials and methods

### Cell lines and culture conditions

HeLa (ACC57) cell lines were purchased from DSMZ and used in experiments between passages 3 to 35. SUDHL-4 and HT were purchased from DSMZ. HEK293TT, PLC/PRF/5, HUH-7, NCI-H226, NCI-H1299, SW620 and HCT116 (p53 +/+, −/−) were a respective gift from Tanveer Ahmad, Zuzana Macek-Jilkova, Morgane Couvet, Hichem Mertani, and Simon Lebaron. HeLa and NCIH226 cell lines were cultured in RPMI Glutamax supplemented with 10% FBS, 1 mM pyruvate sodium, 1% minimum essential medium non-essential amino acids (MEM NEAA) and 100 µg/mL penicillin/streptomycin. HEK293TT, PLC/PRF/5, and HCT116 cell lines were cultured in DMEM Glutamax supplemented with 10% FBS, and 100 µg/mL penicillin/streptomycin. SUDHL-4 and HT cell lines were cultured in RPMI Glutamax supplemented with 20% FBS, 1mM pyruvate sodium, 1% minimum essential medium non-essential amino acids (MEM NEAA) and 100 µg/mL penicillin/streptomycin. All cells were grown at 37 °C, 5% CO_2_. Cell culture reagents were purchased from Gibco. MycoAlert Assay (Lonza) was performed regularly to verify the absence of Mycoplasma.

### Antibodies

The following primary antibodies were used for western blot: CYCLON/CCDC86 (Atlas Antibodies #HPA 04111), GAPDH (Cell Signaling #97166), P53 (Santa Cruz #sc-126), Histone H3 (Milipore #06-755), HSP90b (Genetex – GTX101448) and immunofluorescence: NPM1 (Santa Cruz #sc-271737), Fibrillarin (Abcam, ab4566), Nucleolin (Cell Signaling Technology #14574), Ki-67 (Transduction Laboratories #k72820),

### Activation frequency in cancers and survival analyses

Cancer and non-tumor tissue gene expression data were obtained from GTEX, NCBI SRA and TCGA databases. CYCLON activation threshold was defined as median normal tissue expression plus two standard deviations. Then, the percentage of cancer patients above that threshold (in both negative or positive way) was calculated. Using the same databases, survival was estimated using the Kaplan-Meier method for low and high CYCLON expression groups defined respectively as lower and upper quartile.

### Nucleus and cytosol cell fragmentation

The nuclei of 10 million cells were isolated using a hypotonic nuclei buffer (60 mM KCl, 15 mM NaCl, 5 mM MgCl2, 0.1 mM EGTA pH 8, 15 mM Tris pH 8, 0.3 M sucrose, 1 mM DTT, 1X complete protease inhibitor cocktail, 1X PIC3 phosphatase inhibitor cocktail (Sigma Aldrich), 0.1U/mL of RiboLock (Thermo Fisher) and 0.2% NP40). After a 30 minutes incubation on ice, cells were disrupted 100X using a bounce-cell homogenizer. Nuclei were separated from cytosol by centrifugation (1400 rpm, 5 min, 4 °C). Nuclei integrity was verified using trypan blue staining under an inverted light microscope. Nuclei samples were sonicated at 100% intensity for 1 minute before protein quantification. 10 µg of proteins were resolved by electrophoresis using the procedure described below.

### Immunofluorescence and automatic analysis of nucleoli shape

1 million of suspension cells (DLBCL) were resuspended in PBS and plated in 12 mm glass coverslips for 1h. Adherent cells at 70-90% of confluency plated in 12mm glass coverslips or suspension cells were fixed for 5 minutes in 4% paraformaldehyde (Electron Microscopy Bioscience), permeabilized for 5 minutes with 0.5% (v/v) Triton X-100 (Sigma) in PBS and blocked with 10% Bovine Serum Albumin (BSA, Sigma) in 0.1% Tween PBS (10% BSA-PBST) for 1 hour. Cells were incubated with primary antibodies described above (1/100 dilution in 3% BSA-PBST) overnight at 4°C. then, 3 washes in 3% BSA-PBST were performed, and cells were incubated with goat anti-mouse or anti-rabbit secondary antibody coupled with AlexaFluor 546 or 488 (1:500, Invitrogen) for 1 hour at room temperature and washed 3 times in 3% BSA-PBST. Finally, Hoechst (Thermo Fisher) staining was performed for 3 minutes at 1:1000 dilution in PBS, and coverslips were mounted in a cover slide using Mowiol solution (1:1000, EMD Milipore). Adherent cells images were taken using the Zeiss AxioImager M2 (Zeiss) using the Plan-Apochromat 63×/1,4 Oil lens. Suspension cells microscopy was performed on a confocal microscope (LSM 710, Zeiss) equipped with a 63X/1.4 oil objective using the following settings: laser emission/detection (DAPI 405/405–491 nm; GFP 488/494–550 nm), pinhole 0.7 µM, pixel dwell 3.14 µsec, speed 6, average 4. Images were constructed using ImageJ or ZEN Blue software (Zeiss). Segmentation of nucleus (Hoechst staining) and nucleoli (NPM1 staining) from the background was done individually using the Ilastik software (version 1.4.0). Segmentation probabilities for all raw images uploaded to CellProfiler (version 4.2.6) to obtain nuclei and nucleolar counts and their area. Segmentation outputs were visually inspected before quantification.

### Establishment of Inducible CYCLON knock-down cell lines

shRNA sequences targeting CYCLON (shCyA: CCAGCGTCAGCAAGACCTACA and shCyB: GTCGTCCAAGTGATCCGAAAC or control (shCtrl: CCTAAGGTTAAGTCGCCCTCG) were designed using the DSIR tool (http://biodev.cea.fr/DSIR/DSIR.html). Oligos were annealed and ligated by T4 ligase with the pLKO Tet-On inducible plasmid pre-digested with AgeI/EcoRI (27). Lentiviral particles were produced by the ANIRA platform (ENS Lyon). HeLa, HCT116, SW620, NCI-H1299, NCI-H226, PLC/PRF/5, HUH-7, SUDHL-4 and HT cell lines were transduced with lentiviral particles at multiplicity of infection (MOI) of 10. Viral particles were washed after 24 hours. 1 μg/mL of puromycin was added at day 3 post-infection. Induction was performed with doxycycline (Sigma) at 1 µg/mL during at least 72h. Knockdown efficiency was assessed by Western Blot and qPCR.

### Western blot

Proteins were extracted from 0.5 million cells using RIPA buffer (150 mM NaCl, 50 mM Tris-HCl pH 8.0, 0.5% sodium deoxycholate, 0.1% sodium dodecyl sulfate, 1% NP-40). Lysates were incubated 30 minutes at 4 °C under agitation. Samples were then water-sonicated with VibraCell sonicator (30 seconds ON/OFF for 30 minutes per sample). Then, samples were centrifuged 10 minutes at 16 000g and 4°C to clear insoluble material and quantified with Bradford protein assay. 30 µg of each sample were loaded on a gradient gel NuPAGE 4-12% Bis-Tris (Invitrogen) and run in MES buffer (Invitrogen). The proteins were transferred on a nitrocellulose membrane for 1 hour at 100V at 4°C in Tris Glycine 10% ethanol. Membranes were blocked in either 10% milk or BSA in 0.1% PBS-Tween (PBST) for 1 hour, and incubated overnight with primary antibodies (1/1000 dilution in 3% milk or BSA PBST) in under constant rotation at 4°C. The membranes were washed three times in PBST, incubated with the following secondary antibody: goat anti-rabbit IgG peroxidase 1/10 000 (Merck, #A9169-2ML,) or rabbit anti-mouse IgG peroxidase 1/5000 (Agilent #P026002-2) for 1 hour at room temperature, washed three times and revealed with Luminol/peroxidase mix Clarity Western ECL (Biorad). Acquisitions were realized using Vilber imaging system.

### RNA extraction & RT-qPCR

Total RNA was extracted from 1 million cells with NucleoSpin RNA extraction kit (Macherey-Nagel) according to manufacturer’s instructions. Reverse transcription was carried out following RT SuperscriptTM III kit (Invitrogen RT) from 1μg extracted RNA. Target genes were amplified using the following PCR primers (CYCLONforw: GAGGAGGCACCAAAGTGTTCT, CYCLONrev: AGCTCCAGTAGTGGCTGAGAG, HSPC3forw: ATGGAAGAGAGCAAGGCAAA, HSPC3rev: AATGCAGCAAGGTGAAGACA, GAPDHforw: CCACTCCTCCACCTTTGAC, GAPDHrev: ACCCTGTTGCTGTAGCCA, 45/47Sforw: GAACGGTGGTGTGTCGTTC, 45/47Srev: GCGTCTCGTCTCGTCTCACT) and quantified by qPCR on a thermocycler CF384 Touch Real-Time PCR (Bio-Rad). Ct values (cycle threshold) were analyzed with CFX ManagerTM Software (Bio-Rad) and normalized using geometric mean of GAPDH and HSPC3 values.

### RNA sequencing

Total RNA was extracted and purified according to the procedure mentioned in the RNA extraction & qPCR section. Purity of RNA was monitored by nanodrop analysis. RNA paired-ended sequencing and stranded library were performed by BGI genomics sequencing platform from 500 ng total RNA in 10 µL of ultrapure water. Sequencing depth is 30 million reads per sample. Reads were quality-checked using FastQC. Reads were then aligned to *H. sapiens* genome build hg38 using STAR 2.7.1a (28). Raw read counts for each gene were calculated using HTSeqCount 0.11.2 using defaults parameters (29). Read count data was normalized using the R packages SARTools 1.7.4 (30) and DESeq2 1.32.0 (31). Analyses were performed with R release 4.1.0. DESeq2 normalized read counts were used to identify differentially expressed genes with an adjusted p-value < 0.05 and a fold change below –2 or above 2.

### Gene set enrichment analysis

Normalized counts from all CYCLON-silenced conditions (shA/shB) were analyzed versus the shCtrl in GSEA v4.2.1 (32) using the Hallmarks, C2, C5, and C6 gene sets with more than 15 and less than 500 genes. Gene sets enriched with a fold discovery rate (FDR) q-value lower than 5% were visualized using Cytoscape Enrichment Map plugging (3.9.1).

### Live-imaging

HeLa cells were plated in a labtek and led adhere overnight. Phase contrast images were taken every 30 minutes for at least 72 hours at 40X objective at the Dynamic microscope (Zeiss) with the Evolve 512 – Monochrome 16-bit camera. Mitosis and interphase duration were measured manually.

### Evaluation of 28S/18S rRNA ratio

The RNA Nano (Agilent) chip was prepared following the manufacturer instructions. RNA was denatured by heating 10 minutes at 95 °C and 1 μL was loaded per well. 28S and 18S peaks were identified using the associated software.

### Northern Blot

RNA was extracted from 5-10 million HeLa cells using TRIzol reagent (Invitrogen) according to the manufacturer’s instructions. Total RNAs were resuspended in deionized formamide, denatured at 95°C for 2 minutes and then resolved by electrophoresis as described before (33). Transfer was performed on a membrane. Membranes were incubated overnight in the hybridization buffer containing the probes. Membranes were washed four times and exposed on ChemiDoc. Each rRNA intermediate was normalized on 45/47S pre-rRNA.

### Polysome profiling

10 million HeLa cells in culture were treated with 25 μg/mL of emetine (Sigma-Aldrich) for 15 minutes and lysed in 10mM Tris-HCL pH7.5, 5 mM MgCl2, 100 mM KCl, 1% Triton X-100, 2 mM DTT, 1 U/μL RNAseOUT and 2× cOmplete EDTAFree protease inhibitors. Lysates were centrifuged at 1300 × g for 10 minutes to pellet nuclei. Supernatants corresponding to cytoplasmic fractions were then loaded on 10–50% sucrose gradients poured using the Gradient Master (Serlabo Technologies) and centrifuged at 210.000 × g for 155 minutes at 4 °C. Fractions were collected using the TELEDYNE ISCO collector while concomitantly acquiring corresponding 254 nm absorbance.

### Protein synthesis assay

HeLa cells were plated in 96-well plates overnight and were rinsed with PBS and incubated for 1 hour at 37 °C with 100 μM HPG in methionine-free media (Gibco). The negative control didn’t include HPG, while the positive control 70 µM of cycloheximide. Cells were then washed once with PBS and fixed for 15 minutes in 3,7 % of FA in PBS. Then, the HPG/Alexa Fluor 488 Click-it reaction was performed following the manufacturer’s protocol (Thermo Fisher Scientific #10428). Alexa Fluor 488 mean fluorescence intensities (HPG) and Hoechst 350 mean fluorescence intensities per cell were determined using the ClarioStarPlus automatic plate reader at top detection and spiral scan. HPG values were normalized to the Hoechst values.

### Fluorescent cross-linking and analysis of cDNAs (fCRAC)

The fCRAC experimental and analytical workflow is described in detail in a methodological article by Mikolajczyk, Robertson, Garcia-Sandoval et al. (pending BioRxiv DOI). Here, HeLa cells expressing CYCLON-6xHistidine-4xAlanine-FLAG, with GFP as screening marker, were used for CYCLON fCRAC. HEK293TT cells expressing the SARS-CoV-2 N-Protein and wild-type HeLa cells were included as positive and negative controls, respectively. For each condition, 30 million cells were UV-crosslinked on ice at 254 nm using 400 mJ/cm2. Samples were then processed by tandem purification, 3’ and 5’ IRDye DNA linker ligation, electrophoretic size selection of RNA-protein complexes, reverse transcription, DNA library preparation, and Illumina sequencing. Libraries were sequenced by the Edinburgh Clinical Research Facility on a Nextseq 2000 platform, with 50% PhiX Control v3 (#FC-110-3001) spiked to improve cluster resolution, generating approximately 100 million single-end reads

### Transmission Electron Microscopy (TEM)

HeLa, HCT116, PLC and H226 cells were cultured in 6-wells (Greiner) and fixed using a 1:1 v/v solution of glutaraldehyde 4% and solution of sodium cacodylate 0.2 M pH 7.4 at 4 °C. Cells were washed three times in cacodylate buffer, stained with 2% of OsO4 for 1 hour at 4°C, and dehydrated in series of ethanol. Cells were scraped, included in an Epon epoxy resin, and polymerized at 60°C for 72 hours. Thin sections were cut on a UC7 ultramicrotome (Leica), mounted on carbonic grids and negatively stained with uranyl acetate. Sections were examined with a Jeol 1400JEM 120 kV transmission electron microscope, equipped with a Gatan Orius 600 camera on wide field position and Digital Micrograph software v1.7 (Gatan Inc).

### xCELLigence

HeLa cells plated in 16-well E-plates (Agilent) at a concentration of 5,000 cells/well and let equilibrate for 1 hour. The RTCA program was set to make impedance readings every 5 minutes for 5 hours (adherence) and every 10 minutes for 72 hours (proliferation). Impedance values were normalized to the 5-hour value.

### 3D tumor spheroid assay

1,000 HeLa cells were plated in 96-well in low-binding plate Nunclon Sphera (Thermo Fisher) and centrifuged at 1400 rpm for 5 minutes. Images of spheroids at day 3 and 7 were taken using the Eclipse TS2 light microscope at 40X (Nikon). Quantification of the nucleolar area/nuclear area ratio (to reduce variation due to differences in cell size) was performed in ImageJ software using a custom macro.

### Colony assay

After induction, 50 cells per well were plated in a 24-well plate. The medium was changed every 3-4 days. Cells were then fixed using glutaraldehyde 5% and crystal violet 0.05% solution during 1 hour at room temperature (RT). Wells were cleaned several times using PBS and then let dried. Colonies were counted manually under the criteria that a colony consists of at least 50 cells.

### Cell cycle analysis

1 million cells were harvested and washed one time in PBS. The pellet was resuspended in 100µL PBS and fixed with 900µL of cold ethanol (70%) added drop by drop under constant agitation of the sample collection tube. After 30 minutes, the samples were stored at –20°C for some days. On the day of the analysis, samples slowly thawed at 4°C and then centrifuged (2000 RPM, 10 minutes). The pellet was resuspended in 1mL of PBS in order to complete rehydration of the fixed samples during 4h at 4°C. After rehydration, cells were collected by centrifugation (2000 RPM, 5min) and resuspended in 1mL PBS, 10µg/mL of RNAse A (Sigma Aldrich) and 10 µg/mL propidium iodide (Sigma Aldrich). Samples were incubated at room temperature in the dark during 30 minutes before being analyzed by flow cytometry (BD FACSymphony™ A5 Cell Analyzer). Data were analyzed using ModFit LT software.

### Cell death assay

Cell death was determined by dual Annexin V-FITC (BD Biosciences) and propidium iodide (Sigma Aldrich) staining. First, 0,1 µg/mL Annexin V was diluted in Binding buffer solution (0,1 M HEPES pH7.4, 1.4 M NaCl, 25 mM CaCl_2_). Next, cells were harvested and washed with PBS and stained in 100µL of Annexin V staining mix at room temperature for 10 minutes in the dark. Then, 100 µL of 1X Binding Buffer was added. Finally, 5µg/mL propidium iodide is added before flow cytometry analysis (10 000 events analyzed per condition) at the Attune NxT (Thermo Fisher). Live cells and apoptotic cells were gated in order to get the number of events for each population. Cell viability was determined by the ratio of the number of alive cells events (AnV-/IP-) and that of apoptotic cells events (AnV+/IP-, AnV+/IP+) compared to the total number of events.

### Statistics

All statistics analysis were performed in GraphPad Prism 10.1.2. Data are presented as mean +/− standard deviation. The number of replicates, as well as the statistical tests used for the different comparisons between experimental conditions, are detailed in the figure legends.

## Results

### CYCLON is overexpressed and nucleolar localized across multiple cancer types

CYCLON was defined as a “common essential” gene in DepMap database, based on the large number of cancer cell lines being dependent on CYCLON for their survival. To further assess the general involvement of CYCLON in cancer progression, its expression was compared between TCGA tumor samples and normal tissue profiles from GTEx. Overexpression was defined as expression levels exceeding the median value in normal tissues by more than two standard deviations. Strikingly, CYCLON was upregulated in a large majority of cancer types (23/27) with overexpression observed in 10% to 90% of cases **(Fig. 1A)**. In particular, CYCLON expression was associated with adverse prognosis in Diffuse Large B-cell Lymphoma (DLBCL, as previously reported (7, 8) (**Fig. 1B**, up) and showed significant prognosis impact in hepatocellular carcinoma (LIHC) (**Fig. 1B**, down). CYCLON protein expression was assessed in a panel of cancer cell lines, including HeLa cells, DLBCL, LIHC, LUSC (lung adenocarcinoma), as well as colon adenocarcinomas (COAD). CYCLON levels showed marked variability across cell lines, including within the same subtype **(Sup. Fig. 1A)**. However, based on the information available, this heterogeneity could not be associated with any known cellular characteristics, such as genetics or proliferation rate. We next determined whether CYCLON protein expression levels correlated with cellular dependency on CYCLON for survival using the DepMap database. No correlation was observed between these two parameters **(Sup. Fig. 1B)**. These findings indicate that, although frequently overexpressed in cancer, CYCLON’s expression level is not predictive of dependency across cell lines, suggesting that functional reliance is context-dependent rather than expression-driven.

**Figure 1.**
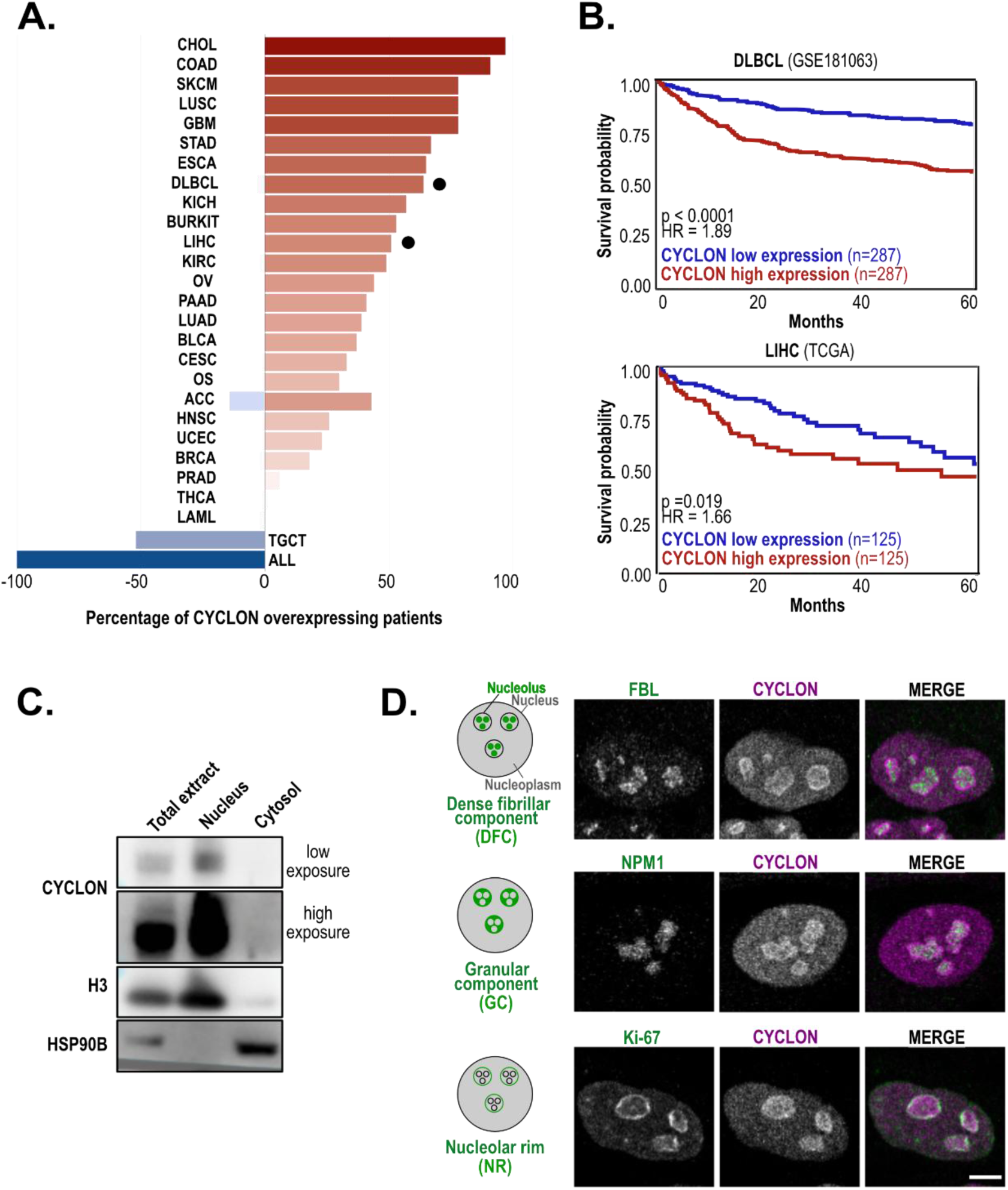
CYCLON expression and localization in cancer. **A.** Percentage of CYCLON overexpression in cancer patients (TCGA database). Cancer types are indicated according to TCGA Study Abbreviations. **B.** Kaplan-Meier survival curves for CYCLON high (upper tercile) and low (lower tercile) expression in DLBCL (GSE181063) and LIHC (TCGA) datasets. HR: Hazard Ration, p value derived from a log rank test **C.** Western blot analysis of CYCLON expression in Hela cells nuclear and cytoplasmic fractions. Histone 3 (H3) and HSP90B were used respectively as nuclear and cytosolic markers. **D.** Immunofluorescence analysis of endogenous CYCLON co-localized with markers of specific nucleolar regions (indicated in the left scheme) in HeLa cells. FBL, Fibrillarin. All images were taken in a confocal microscopy at 63X, bar = 5 µm. All experiments were done in triplicate.

Immunofluorescence analyses confirmed that CYCLON is a nuclear protein, as previously reported (10, 13). Localization was observed in both the nucleolus and the nucleoplasm, with variable distribution between these compartments among different cell lines **(Sup. Fig. 1C)**. Cell fractionation experiments confirmed an exclusive nuclear localization, as CYCLON was absent from the cytosol after isolating nuclear and cytosolic compartments. **(Fig. 1C)**. Within the nucleolus, CYCLON is preferentially localized to the granular component (GC) and the nucleolar rim (NR) in HeLa cells, as evidenced by its colocalization with NPM1 and Ki-67, respectively **(Fig. 1D).** Localization to the granular component suggests a role in ribosome maturation, while the nucleolar rim is regarded as a transitional zone enriched in proteins involved in stress sensing, nucleolar structural maintenance and interaction with heterochromatin (34).

Taken together, these data suggest that CYCLON could be an underestimated player in cancer cell biology, exhibiting a shared interfacial localization between the nucleoplasm and the nucleolus. This pattern suggests a potential role in nucleolar functions and proliferation-associated pathways.

### CYCLON silencing affects nucleolar morphology

To identify CYCLON functions, we developed an inducible knock-down (KD) model through lentiviral transduction of CYCLON-targeting short-hairpin RNA (shRNA) in HeLa cells (27). After 72h of induction, a maximal reduction of approximately 80% in CYCLON mRNA and protein levels was obtained using two distinct target sequences (shCyA and shCyB) compared to control cells transduced with a control shRNA (afterwards referred as shCtrl) **(Sup. Fig. 2A-B).**

**Figure 2.**
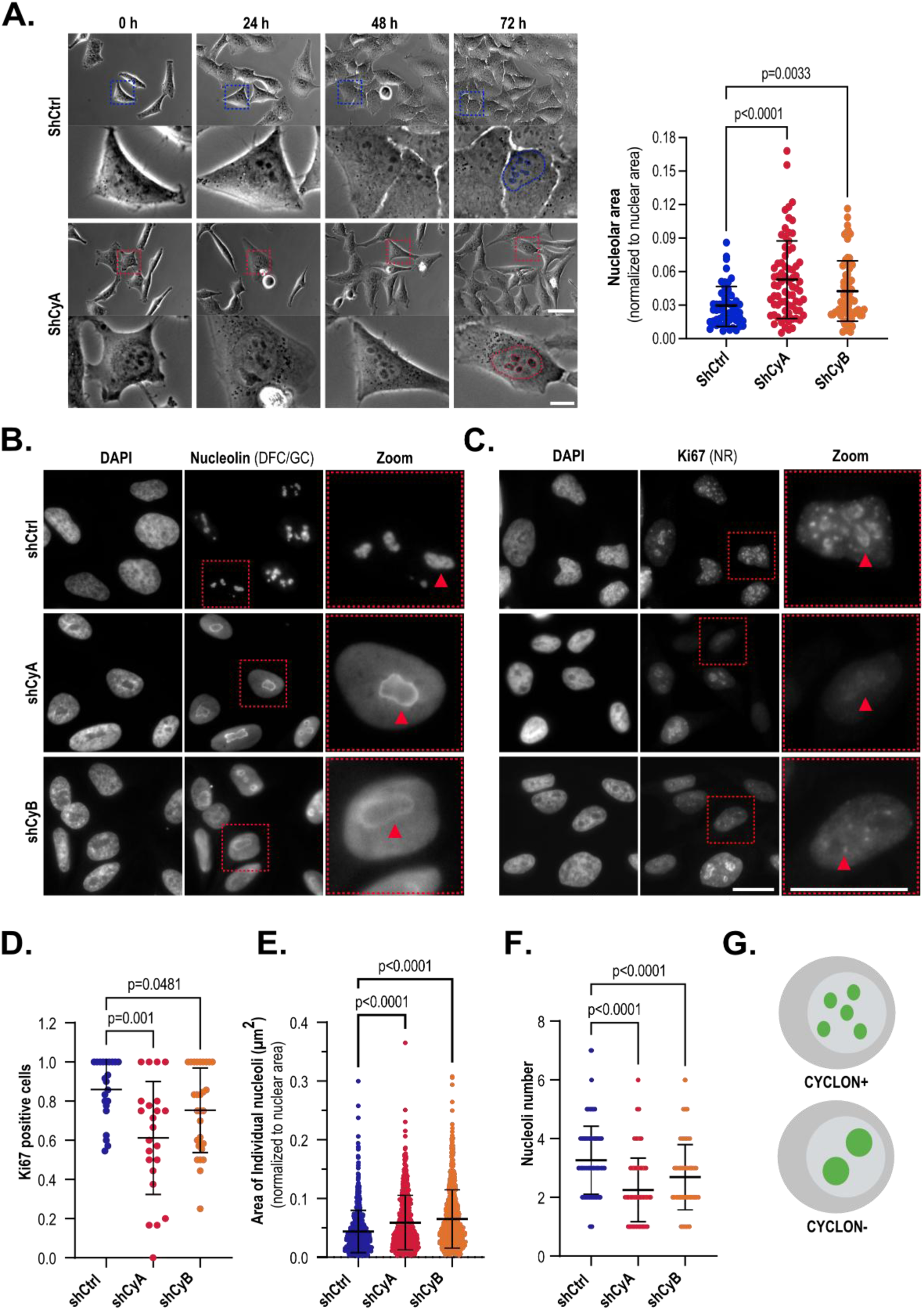
CYCLON affects nucleolar morphology in HeLa cells. **A.** (left) Phase-contrast live-images in cells transduced with a control (shCtrl) and CYCLON targeting shRNA (shCyA and shCyB) after 72h doxycycline induction and additional 72h of imaging. Scale bar = 50 µm, 10 µm (zoom). N = 12 positions with 8-10 cells each, three independent experiments. The squares indicate the same cell followed through the time-lapse and shown in zoomed view. (right) Manual quantification of nucleolar area (normalized to nuclear area) after 72h. p values were derived from Kruskal-Wallis test followed by uncorrected Dunn’s test. **B-C.** Representative immunofluorescence images of Nucleolin (B) and Ki-67(C) and DAPI nuclear staining, in cells transduced with a control (shCtrl) and CYCLON targeting shRNA (shCyA and shCyB) for 96h. Scale bar 20 µm, n = 30 images with 5-10 cells each, three independent experiments. The red arrows indicate morphological differences between shCy and shCtrl cells, while the dotted squares highlight the zoomed-in regions. **D.** Quantification of Ki-67 positive cells from panel B images. p values were derived Brown-Forsythe ANOVA followed by un-paired t-test with Welch-s correction. **E-F.** Automated quantification of NPM1 fluorescence signal to evaluate nucleoli area (E) and number (F), n = 250 cells, three independent experiments. p values were derived from a Kruskal-Wallis test followed by uncorrected Dunn’s test). **G.** Scheme representing nucleoli in CYCLON-expressing (CYCLON+) or CYCLON-silenced (CYCLON-) cells

A transcriptomic analysis performed 96h post-induction revealed a modest transcriptional response upon CYCLON depletion, with 74 genes (36 down, 28 up) showing altered expression (FC<0.5 or >2, p<0.05) **(Sup. Fig. 2C)**. Enrichment map analysis of the significant GSEA gene sets (FDR q-value < 0.05) identified three major clusters of biological functions: rRNA biogenesis, mitochondrial translation, and immune signatures **(Sup. Fig. 2D)**. Other nucleolar proteins, such as nucleolin in HeLa cells (35), also showed modest transcriptome-wide effects, these findings supporting a role for CYCLON in ribosome production.

Our inducible knockdown strategy allowed to monitor the effects of CYCLON loss over a 3-days period required to observe a maximal reduction in protein levels. Evolution of the morphological phenotypes of CYCLON KD cells was first inspected by phase-contrast live-imaging on a cell-by-cell basis. CYCLON loss induced a significant enlargement in nucleoli size after 72 hours of induction, suggesting perturbations of nucleolus morphology (**Fig. 2A).**

To further investigate morphological changes within the nucleolus, its structural integrity was assessed by fluorescence microscopy using specific markers for each nucleolar sub-compartment. As markers, we included nucleolin, fibrillarin (36, 37), NPM1 (13, 38, 39), and Ki-6, which localize respectively at the dense fibrillar component (DFC), granular component (GC), and nucleolar rim (NR). Each has been reported to interact with CYCLON in high-throughput screens (Biogrid, MINT and Intact databases). Following CYCLON depletion, nucleolin was delocalized from the GC to the nucleolar rim and nucleoplasm **(Fig. 2B).** Similar nucleolin relocation has been reported upon nucleolar stress (40–42). Notably, Ki-67 was both relocated to the nucleoplasm **(Fig. 2C)** and significantly reduced in CYCLON-depleted cells (**Fig. 2D**). While the localization of fibrillarin **(Sup. Fig. 2E)** and NPM1 **(Sup. Fig. 2F)** appeared unchanged, the regions occupied by the DFC (decorated by fibrillarin) and the GC (decorated by NPM1) were visibly enlarged. Changes in nucleolar morphology were automatically quantified based on NPM1 staining. This confirmed the increase in nucleolar area **(Fig. 2E)** and revealed a decrease in nucleolar number **(Fig. 2F** schematized in **Fig. 2G)**. Together, these findings indicate that CYCLON silencing disrupts nucleolar organization, leading to structural remodeling and features consistent with nucleolar stress.

### CYCLON affects ribosome biogenesis and protein synthesis by inhibiting pre-rRNA processing

To assess whether the morphological changes observed upon CYCLON loss are linked to defects in nucleolar functions and ribosome biogenesis, the 28S/18S rRNA ratio was measured from Bioanalyzer electropherograms. This ratio was shown to be significantly decreased upon CYCLON depletion (**Fig. 3A**). Polysome profiles confirmed that CYCLON depletion reduces 60S subunits relative to 40S and leads to a modest decrease in translation-competent polysomes, consistent with impaired ribosome assembly and reduced translational output (**Fig. 3B**). This was further confirmed by HPG Click-IT assays, which reveal a marked decrease in global protein synthesis, comparable to that observed with the translation inhibitor cycloheximide (∼4-fold reduction) (**Fig. 3C**).

**Figure 3.**
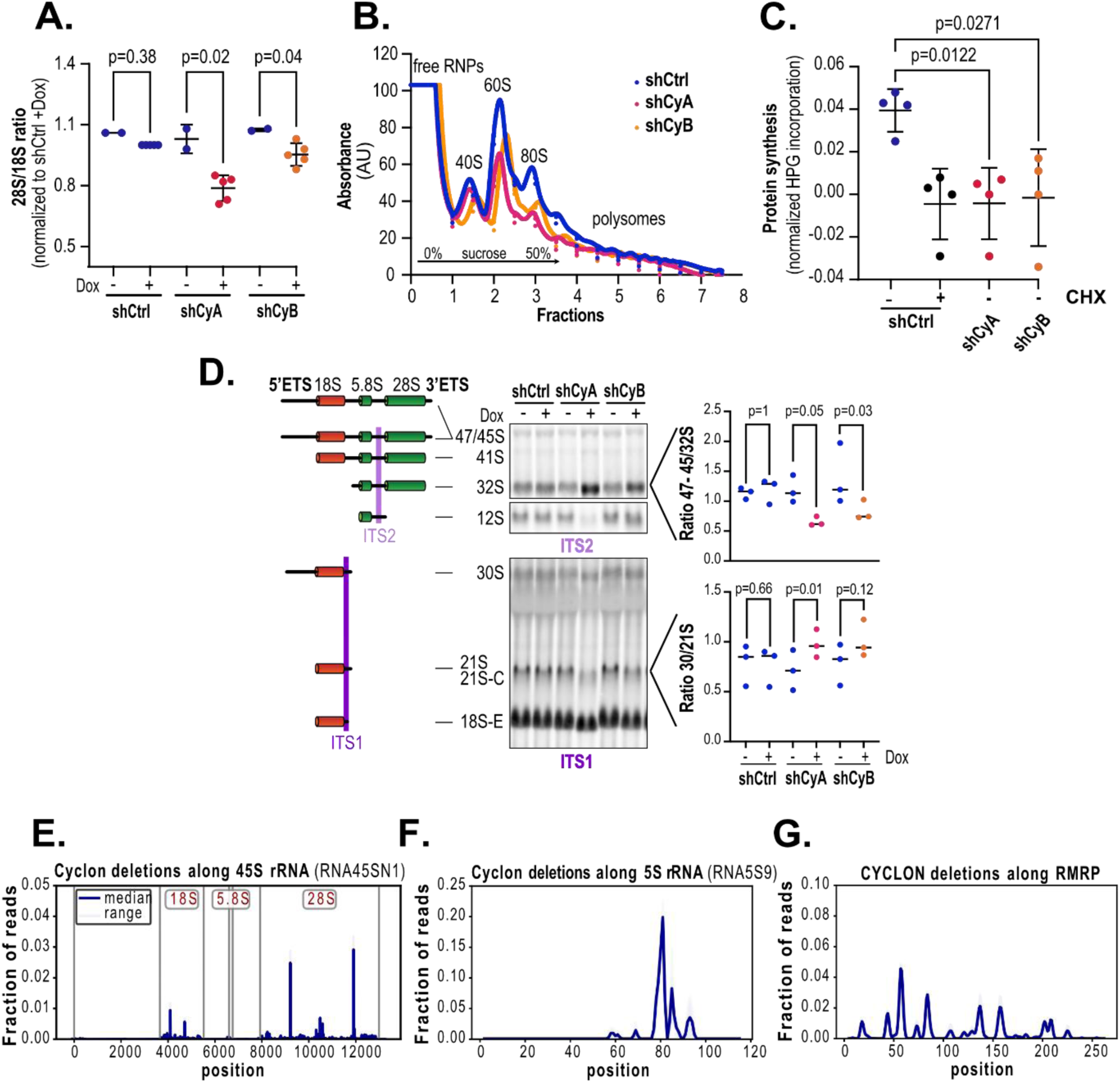
CYCLON plays a role in ribosome biogenesis in HeLa cells. **A.** 28S/18S rRNA ratio determined by bioanalyzer in cells transduced with a control (shCtrl) and CYCLON targeting shRNA (shCyA and shCyB) after 96h doxycycline induction, n = 2 for Dox (−), and 5 for Dox (+). **B.** Representative polysome profile in cells transduced with a control (shCtrl) and CYCLON targeting shRNA (shCyA and shCyB) after 96h doxycycline induction, n = 3. **C.** Protein synthesis measured by HPG incorporation in cells transduced with a control (shCtrl) and CYCLON targeting shRNA (shCyA et shCyB) after a 96h doxycycline induction. Cycloheximide (CHX+) was used as a positive control for protein synthesis inhibition, n = 4. **D.** (left) Scheme of intermediate rRNA species detected using ITS1 and ITS2 probes, (middle) representative Northern Blot analysis of cells expressing a control (shCtrl), CYCLON targeting shRNA (shCyA and shCyB) using ITS1 and ITS2, (n=3), quantification of ratio between rRNA intermediates derived from Northern blot analyses as indicated, n = 3, p values are derived from a Friedman ANOVA followed by uncorrected Dunn’s test. **E-G.** fCRAC mapping of deletions detected by sequencing CYCLON UV-crosslinking on 45S (E) and 5S rDNA (F), as well as RMRP loci (G).

Levels of ribosomal RNA precursors (pre-rRNAs) were assessed to determine whether this deficit in 60S subunits reflects impaired ribosome synthesis. The level of the 47S primary transcript was assessed by RT-qPCR performed against a region in the 5’ETS of the 45S rRNA as previously described (43). his suggested that pre-rRNA transcription is unaltered (**Sup. Fig. 3A**), consistent with CYCLON absence from the inner nucleolar regions, FC and DFC. Northern blot analyses were performed using two specific probes targeting the ITS1 and ITS2 regions to precisely identify which steps of rRNA processing were impacted upon CYCLON depletion. Both CYCLON-targeting hairpins (shCyA and shCyB) consistently caused accumulation of the 32S (60S ribosomal subunit precursor) and decrease in 21S pre-rRNA intermediates (40S ribosomal subunit precursor), as quantified by altered precursor-to-intermediate ratios (47S+45S/32S and 30S/21S) **(Fig. 3B)**. The levels of the 12S rRNA intermediate (**Sup. Fig. 3B**) exhibit variability between CYCLON-targeting sequences suggesting that this change may require a specific dynamic or threshold of CYCLON depletion not met upon shCyB depletion, even though this could not be experimentally detected. Taken together, these data suggest that CYCLON impairs early-to-intermediate steps of pre-rRNA processing affecting both large and small ribosomal subunits maturation, that could be related to the observed changes in nucleolar morphology.

Both CYCLON’s nucleolus localization and role in rRNA processing may be mediated through direct physical contacts with specific regions of the pre-rRNA or other type nucleolar RNAs participating to this process. This hypothesis was tested using fCRAC, a fluorescent-based adaptation of CRAC (Crosslinking and Analysis of cDNAs) analysis (44). This protocol uses UV cross-linking to stabilize direct RNA–protein interactions prior to denaturing purification of bipartite-tagged proteins, enabling the mapping of rRNA-interacting proteins with low background (45–47). fCRAC revealed precise cross-linking sites within the mature 18S and 28S sequences of the pre-rRNA transcripts (**Fig. 3E**), as well as specific interactions with the 5S rRNA (**Fig. 3F**). Of the uniquely mapping RNAs (i.e., excluding the multicopy rRNA species), the highest ranked species was RMRP, the catalytic RNA core of the RNase MRP complex that cleaves the ITS1 at site 2 in humans (44, 48) (**Fig. 3G**). These results confirm CYCLON’s direct rRNA binding and involvement in pre-rRNA processing complexes. In addition, fCRAC revealed that CYCLON makes contact with numerous snoRNAs (**Sup. Fig. 3C**), suggesting wider roles in promoting pre-ribosome assembly and pre-rRNA modification. Notably, the dominant CYCLON targets in fCRAC were tRNAs (**Sup. Fig. 3C**). Potential roles of CYCLON in tRNA maturation lies beyond the scope of this paper, but we speculate that tRNA defects may further reinforce the strong reduction in total protein synthesis seen following CYCLON depletion.

Taken together, these data show that CYCLON acts as a multivalent RNA-binding platform that engages both 40S and 60S maturation pathways and the key processing complex RNase MRP. Loss of these interactions provides a molecular basis for the pre-rRNA processing defects and 60S imbalance observed upon its depletion. These findings therefore identify CYCLON as a regulator of ribosome biogenesis that coordinates early cleavage events with later maturation steps to sustain translational capacity and proteostasis.

### CYCLON loss activates nucleolar stress responses in cancer cell lines

Defects in essential nucleolar proteins involved in pre-rRNA processing and ribosome assembly, particularly incorporation of the 5S RNP, can trigger the nucleolar stress response (49). The nucleolar stress response can proceed through both p53-dependent (through sequestration of the ubiquitin-ligase MDM2 can lead to p53 stabilization, or p53-independent pathways, enabling cells to modulate cell cycle arrest, apoptosis, or adaptive stress programs according to their molecular context (50). We therefore examined whether loss of CYCLON activates this pathway. Ultrastructure analyses of nucleoli were performed by electron microscopy in four cancer cell models of CYCLON knock-down. These consistently revealed enlargement of electron transparent structures (white regions indicated in **Fig. 4A**), which may correspond to nucleolar cavities, usually formed by perinucleolar chromatin projections. These structures are poorly characterized, but they are indicative of nucleolar dysfunction and stress response (51, 52). We then evaluated the impact of CYCLON loss on P53 protein levels in p53 wild-type (HCT-116 and H226) and mutated (HT, SUDHL-4, HUH-7, PLC/PRF/5, SW620). CYCLON depletion led to elevated p53 levels across all cell lines, with a slightly stronger effect observed in the p53 wild-type background **(Fig. 4B-C)**. Stabilization of p53 is consistent with activation of the nucleolar stress response, as previously reported (50), and supports a central role for CYCLON in ribosome biogenesis and the maintenance of nucleolar integrity

**Figure 4.**
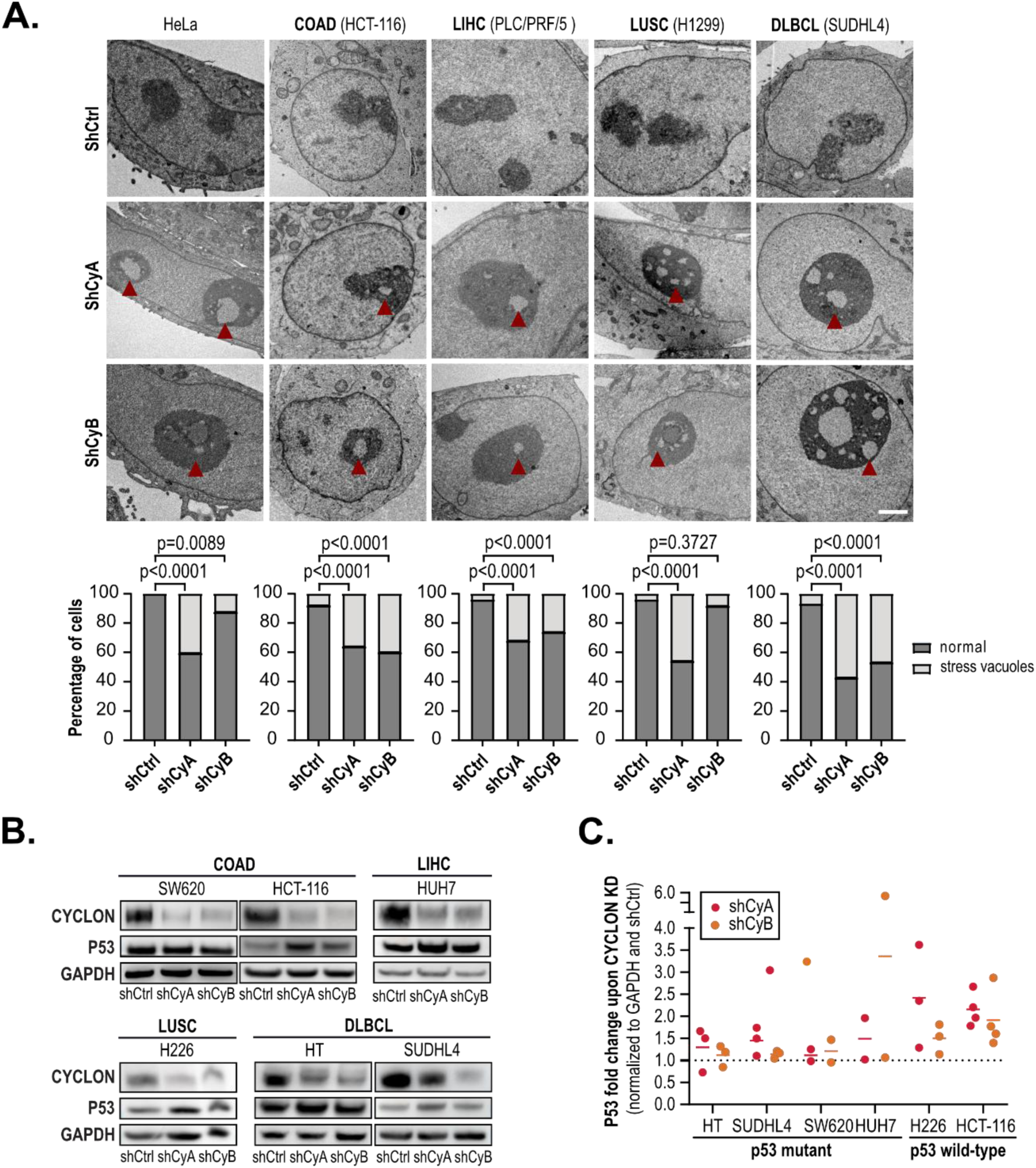
CYCLON loss activates nucleolar stress responses in cancer cell lines. **A.** (up) Transmission electron microscopy images of in cells transduced with a control (shCtrl) and CYCLON targeting shRNA (shCyA et shCyB) after a 96h doxycycline induction in pan-cancer cell lines as indicated, scale bar 2µm, (bottom) quantification of enlargement of white nucleolar sub-compartment, n=40 cells, two independent experiments, p values are derived Fisher exact test. **B.** Western blot analysis of CYCLON and p53 protein expression in cells transduced with a control (shCtrl) and CYCLON targeting shRNA (shCyA et shCyB) after a 96h doxycycline induction in pan-cancer cell lines as indicated. GAPDH levels were used for normalization (n = 3). **C.** Quantification of changes in P53 levels in CYCLON-silenced cells (shCyA and shCyB) normalized to GAPDH and shCtrl cells.

### CYCLON sustains cancer cell proliferation and clonogenic capacity by regulating cell cycle duration

By sustaining the cell’s biosynthetic capacity and enhancing its ability to sense and adapt to stress, CYCLON’s nucleolar functions are likely to promote proliferation, survival and overall fitness of cancer cells. Indeed, we observed in the time-lapse experiment presented in **Fig. 1A**, that CYCLON-silenced cells divided less frequently appeared larger, suggesting a global decrease in cell proliferation. This reduced cell growth was further confirmed in both 2D cultures (followed dynamically by xCELLigence, 17-25% decrease in growth rate) **(Fig. 5A)** and 3D tumor spheroids assay (9-13% decrease) **(Sup. Fig. 4A)**. Consistently, CYCLON expression correlates with that of Ki-67 and PCNA, two established markers of cancer cell proliferation in clinical samples **(Sup. Fig. 4B)**. To assess whether this limited cell growth reflects the ability of cells to divide and increase in number over a short period or true self-renewing and long-term proliferative capacity, clonogenic assays were performed **(Fig. 5B)**. Strikingly, CYCLON loss was associated with a highly significant decreased capacity to form colonies (83-91%), showing its essential role in maintaining cell fitness. This loss of clonogenic capacity was also observed in the vast majority of solid **(Fig. 5C)** and hematological **(Fig. 5D)** cell line models tested, suggesting that this function is conserved across all cell types expressing CYCLON. Interestingly, HCT-116 and H1299 cell lines show the most drastic phenotypes, despite the absence of association between CYCLON expression and prognosis in colon and lung cancer.

**Figure 5.**
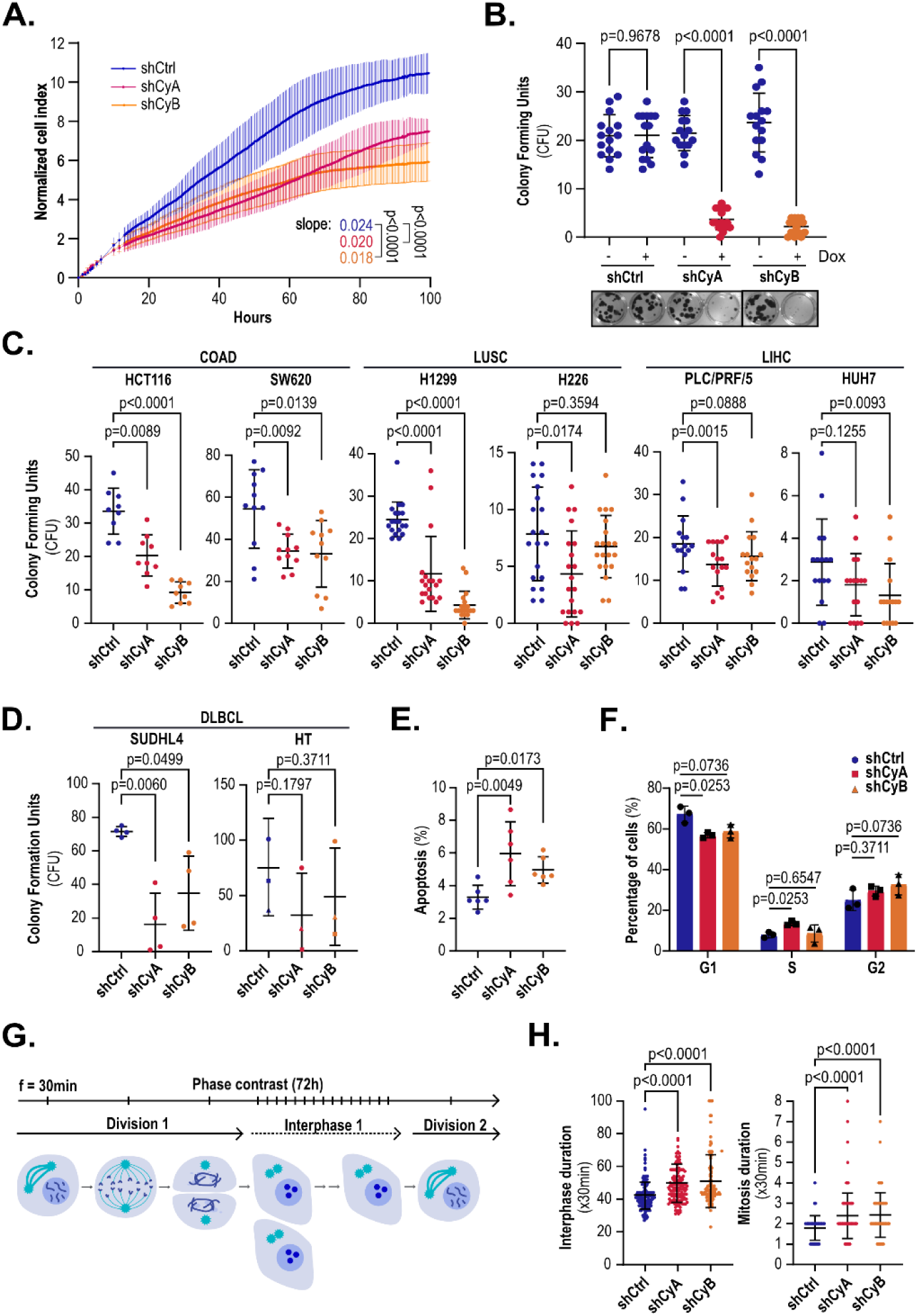
CYCLON-silenced cells present a reduced cell fitness in pan-cancer manner. **A-H.** All experiments were performed in cells transduced with a control (shCtrl) and CYCLON targeting shRNA (shCyA et shCyB) after a 96h doxycycline induction. **A.** Proliferation index over-time measured by xCELLigence assay in HeLa cells, n = 6, three independent experiments. Slopes were calculated from logarithmic transformation of values at the exponential growth phase (from 20 to 60h) and compared using an ANOVA test. **B.** Clonogenic assay in HeLa cells), n = 15, five independent experiments. Representative pictures are displayed in lower panel. **C.** Clonogenic assay in pan-cancer cell line panel, n = 9, three independent experiments. **D.** Clonogenic assay in SUDLH4 and HT DLBCL cells, n = 4, two independent experiments. B-D: for normally distributed cell lines, a parametric test was used (Brown–Forsythe followed by Welch’s t-test), whereas non-normally distributed data were analyzed using a non-parametric approach (Kruskal–Wallis followed by Dunn’s test. **E.** Apoptosis rate in HeLa cells, n=6, p values were derived from a Kruskal-Wallis test followed by uncorrected Dunn’s Test. **F.** Cell cycle analysis performed in HeLa cells, n=6, p values were derived from a Kruskal-Wallis test followed by uncorrected Dunn’s Test. **G.** Schematic representation of time-lapse microscopy analysis of mitosis and interphase duration. Images have been captured every 30 minutes during 72h. **H.** Interphase (left) and mitosis (right) duration in HeLa cells quantified from phrase-contrast live-imaging.

To investigate the biological processes sustaining the reduced proliferation rate observed in these different assays, apoptosis and cell cycle analyses were performed in HeLa cells. CYCLON silencing results in a modest increase in apoptosis (**Fig. 5E**) and a slight G1 arrest, which reaches statistical significance only with the shCyA sequence (**Fig. 5F**). To evaluate whether these effects could be related to nucleolar surveillance pathway and amplified in a p53 wild-type background, we tested p53 wild-type (+/+) and null (−/−) HCT-116 cells. P53 levels and CYCLON-silencing were first validated by western blot (**Sup. Fig. 4C).** We observed that clonogenic capacity is equally affected by CYCLON depletion in both backgrounds (**Sup. Fig. 4D)**. Consistently, we found no association between reduction in clonogenic capacity and p53 status in our pan-cancer cell line panel (**Sup. Fig. 4E).** No significant effect of CYCLON-silencing could be observed on apoptosis (**Sup. Fig. 4F)** or cell cycle (**Sup. Fig. 4G)** of HCT116 cells in neither background. This strongly suggests that the cell-fitness related phenotypes are not enhanced when p53-dependent nucleolar stress response is functional. This data also implies that cell cycle arrest or induction of apoptosis are not the main drivers of CYCLON-induced growth defects.

To investigate this difference in proliferation, live imaging was used to monitor cell division dynamics under inducible CYCLON silencing, tracking the interval between two mitotic events (G0–interphase duration) in single cells (**Fig. 5G**). CYCLON knockdown cells exhibited a significantly prolonged interphase as well as an increased duration of mitosis **(Fig. 5H)**. These results indicate that CYCLON silencing delays cell growth, by extending both interphase and mitosis.

CYCLON therefore emerges as a central determinant of cell fitness, coordinating rRNA processing and ribosome biogenesis to sustain protein synthesis, cell-cycle progression and clonogenic growth. CYCLON’s essential role in sustaining the elevated translational demands of cancer cells underscores its importance for tumor growth and identifies it as a compelling therapeutic vulnerability.

## Discussion

Public data mining revealed that CYCLON is broadly overexpressed across multiple tumor types, where elevated levels can be associated with unfavorable prognosis in distinct subsets, including Liver Hepatocellular Carcinoma (LIHC), Diffuse Large B Cell Lymphoma (DLBCL) (7, 8), nasopharyngeal carcinoma and neuroblastoma (5, 9). Together with its classification as a common essential gene in DepMap database, this indicated that CYCLON was likely involved in fundamental cellular processes. Our integration of phenotypic data across a diverse set of cancer cell lines, revealed that, CYCLON connects the key processes of ribosome biogenesis, nucleolar dynamics and protein synthesis.

CYCLON is enriched in the granular component (GC) and nucleolar rim (NR), and its depletion induced the delocalization of key nucleolar proteins (nucleolin and Ki-67) to the nucleoplasm. Although these changes appear selective, delocalization of these proteins has been reported in several stress conditions such as transcriptional inhibition (25, 42), kinase inhibitors (25), heat-shock and ionizing radiation (40) or knockdown of nucleolar scaffold proteins (41). In addition, CYCLON depletion induced a general increase in nucleolar size and reduction in number. Larger nucleoli are often linked to elevated ribosome biogenesis (53), but our results indicate that nucleolar size alone does not necessarily reflect functional capacity. CYCLON loss impaired pre-rRNA processing and 60S accumulation, but increased nucleolar size, apparently through the formation of large internal vacuoles, previously linked to nucleolar stress (51).

CYCLON shares a putative RNA-binding domain with the yeast ribosome synthesis factor Cgr-1 (12). However, CYCLON also presents a large, N-terminal, intrinsically disordered region (IDR). This suggests a role liquid–liquid phase separation (LLPS), a biophysical mechanism believed to underpin nucleolar organization (54). Loss of this activity may contribute to nucleolar size increase, vacuole formation, and altered nucleolar protein distribution, following CYCLON depletion.

The nucleolus is a robust structure that is not easily altered: high-throughput knock-down of various nucleolar proteins identified only 10% that cause clear disruption of nucleolar morphology (55, 56), including key nucleolar proteins such NPM1 (57, 58). This underscores the central and distinctive role of CYCLON in maintaining nucleolar homeostasis. Besides these morphological changes, CYCLON facilitates nucleolar function in ribosome biogenesis. Loss of CYCLON impaired pre-60S RNA processing, including accumulation of 32S and altered 21S production, and was associated with reduced mature 60S subunits relative to 40S. In vivo crosslinking showed that CYCLON was associated with the mature rRNAs, presumably in the context of pre-ribosomal particles, particularly the 5S rRNA. In agreement with this, a recent Perturb-seq analysis (59) and cryo-EM data (60, 61) indicated that CYCLON may participates in the rotational rearrangement and incorporation of the 5S RNP particle, a key checkpoint in 60S maturation. CYCLON was also associated with RMRP the RNA component of the endonuclease RNase MRP. As specific cleavage in the ITS1 region of the pre-rRNA requires RMRP activity, we speculate that the impaired pre-rRNA processing observed following CYCLON depletion may reflect impaired RMRP association and/or activity.

The inhibition of protein synthesis following CYCLON depletion was very marked, and more important than just a reflection of a reduction in ribosomal subunits. We note that CYCLON was strongly crosslinked to tRNAs in vivo, and speculate that it may also play an, as yet unidentified, role in tRNA maturation.

We revealed the strong impact of CYCLON on clonogenic potential in multiple cancer models, suggesting the nucleolar function is required for long-term self-renewal, proliferative and survival capacities. This was corroborated by shorter-term 2D and 3D proliferation assays (x-CELLigence and spheroids). In previous studies, we showed that CYCLON was increasing tumor growth and reducing response to immunotherapy in lymphoma cells xenografted in mice (7). Collectively, our findings identify CYCLON as a central determinant of cancer cell fitness across tumor types, validated from simple cellular models to *in vivo* systems.

Reduced clonogenic potential and proliferation upon CYCLON loss could be related to the induction of nucleolar stress and the subsequent activation of p53-dependent and independent nucleolar stress responses. Although we observed p53 stabilization upon CYCLON loss in a p53 wild-type background, similar phenotypes were also seen in p53-null cells, suggesting that p53-independent pathway could be related to growth defects observed in this setting (62, 50, 63). Additionally, no alterations were detected in cell cycle progression or apoptosis, classical cellular phenotypes downstream p53 activation (50). Importantly, this finding has clinical relevance, suggesting that CYCLON targeting may remain effective even in p53-deficient cancers, potentially through alternative, yet uncharacterized, stress response pathways Alternatively, CYCLON-induced cell growth defects could also be related to the previous report of mitotic abnormalities, including an increased spindle length and delays in chromosome segregation upon CYCLON loss (5). Here, we identified a general extension of both mitosis and interphase duration which could be linked to impaired protein synthesis, including quantitative and qualitative alterations of ribosomes and their associated translatomes. These alterations would extend cell growth phases and delay division, effects not easily captured by classical relative cell cycle analysis. By facilitating protein synthesis, CYCLON expression can shorten the cell cycle, a typical trait of precursor cells consistent with its normal expression profile in progenitor and proliferating cells. This mechanism is consistent with recent study showing that that reduced cell cycle duration is a hallmark distinguishing cancer-prone from cancer-resistant cells (64), ultimately enhancing cell fitness for oncogenic potential.

Our results also provide further insights into the reciprocal regulation between CYCLON and the proliferation marker Ki-67, another IDR-containing protein enriched at the nucleolar rim. CYCLON silencing was already shown to cause the cytoplasmic accumulation of Ki-67 and nucleolin in NDF-like foci during mitosis, whereas Ki-67 depletion is leading CYCLON delocalization during mitosis (5). Here, we show that CYCLON silencing leads to reduced levels and relocation of Ki-67 during interphase. Although Ki-67 is necessary for maintaining peri-chromosomal architecture and is widely used as a proliferation marker, it is not strictly required for cell viability (65). In contrast, the essential role of CYCLON in coordinating ribosome biogenesis, maintaining nucleolar integrity, and supporting cell cycle progression highlights its broader impact on cellular homeostasis. This raises the question of whether CYCLON could be a more mechanistically informative proliferation marker than Ki-67 in certain contexts. However, specific signals or conditions that CYCLON might sense to influence cellular fitness remain elusive.

This work also highlights the fact that understudied proteins, such as CYCLON, offer opportunities to uncover new aspects of cell biology. Future studies should further dissect its mechanistic contributions to 40S and 60S assembly, explore its interplay with known nucleolar stress pathways, describe its potential sensing activity of the nuclear microenvironment

Collectively, our findings identify CYCLON as a central regulator of nucleolar integrity, ribosome biogenesis, and protein synthesis, thereby sustaining cancer cell fitness across tumor types. By coupling structural organization of the nucleolus with functional control of rRNA processing and translational output, CYCLON emerges as a key determinant of proliferative capacity in cancer cells. It’s essential and pan-cancer nature, together with its impact on clonogenic growth and cell cycle dynamics in both p53-proficient and –deficient contexts, highlights CYCLON as a previously unrecognized vulnerability in tumor biology. These results position CYCLON as both a mechanistic link between nucleolar homeostasis and cellular fitness, and a promising candidate for future biomarker and therapeutic exploration.

## Acknowledgments

This work was supported by the French Ministry of Higher Education, Research and Innovation (MESRI), which provided a PhD scholarship to AGS, as well as AMGEN Innovations (Fond AMGEN France pour la Science et l’Humain) and La Ligue Contre le Cancer (Appel d’Offres Régional, CC-AURA 2022) awarded to AE. NR was supported by a Scottish Government Chief Scientist Office/NHS Education for Scotland Clinical Lectureship (PCL/23/07) and an Academy of Medical Sciences Starter Grant for Clinical Lecturers award (SGCL033\1114).

We thank the Microcell core facility of the Institute for Advanced Biosciences (UGA – Inserm U1209 – CNRS 5309), especially Mylene Pezet and Solenne Dufour, for their assistance with the Attune NxT equipment. This facility belongs to the IBISA-ISdV platform, member of the national infrastructure France-BioImaging supported by the French National Research Agency (ANR-10-INBS-04). We acknowledge the contribution of SFR Santé Lyon-Est (UAR3453 CNRS, US7 Inserm, UCBL) CIQLE facility (a LyMIC member), especially Elisabeth Errazuriz-Cerda and Christel Cassin, for their help in sample preparation and electron microscopy. We recognize the contribution of SFR Biosciences (UAR3444/CNRS, US8/Inserm, ENS de Lyon, UCBL) AniRA facility from the CELPHEDIA Infrastructure (Gisèle Froment, Didier Nègre and Caroline Costa) for lentiviral particles production.

## Author contributions

AGS, SD, NR, SH, FB and VL performed experiments and analyzed data. EM performed bioinformatics and biostatistics analysis. AGS, SD, NR, DT, JJD, OD and AE designed the study. OD, AE coordinated the study. All authors reviewed the manuscript.

## Data availability

Sequencing data are being deposited on GEO database (pending accession number). All other data that support the findings of this study are available from the corresponding author upon request.

---

**Supplemental Figure 1.**
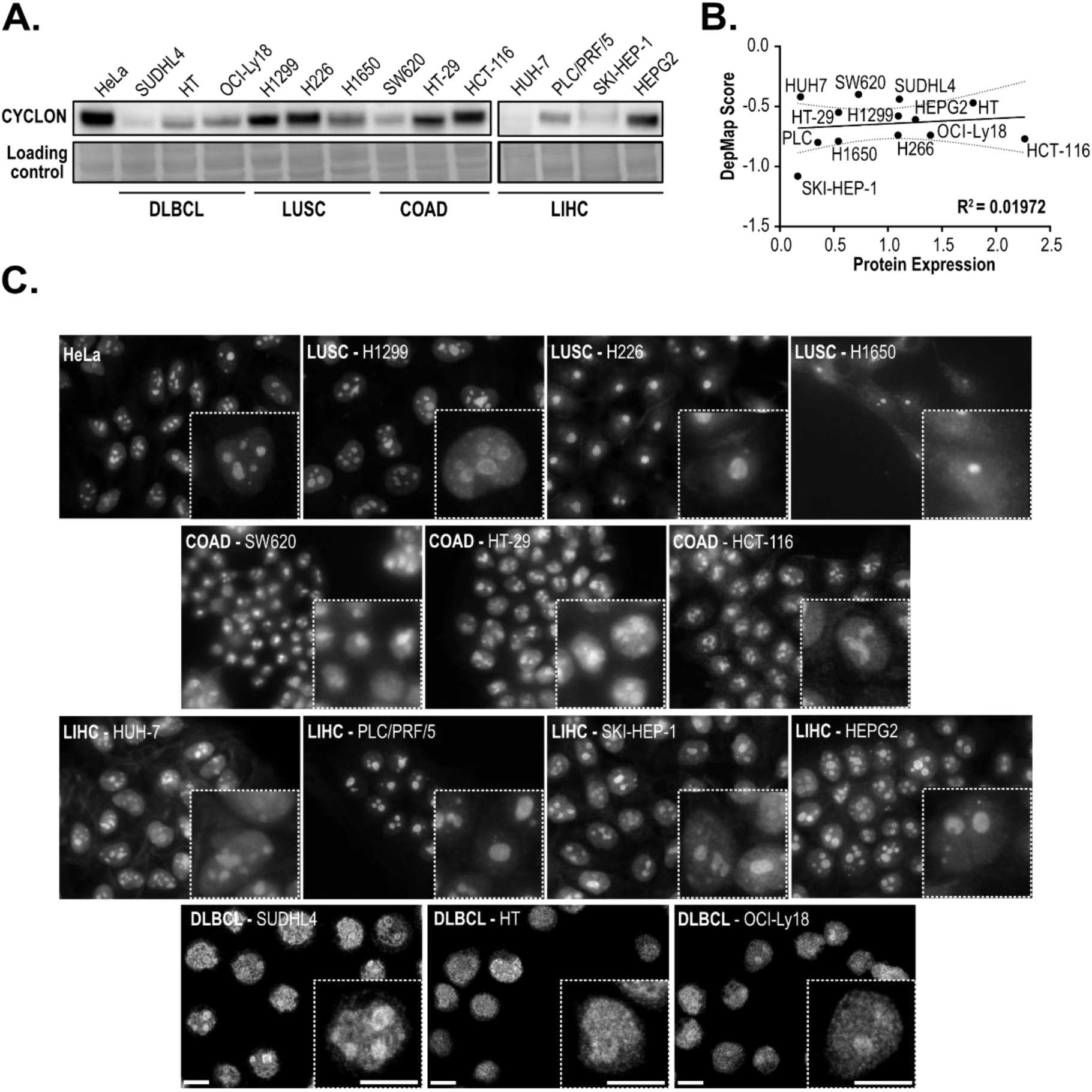
CYCLON expression and localization in cancer. **A.** Western blot analysis of CYCLON protein levels in a pan-cancer cell line panel. Ponceau staining was used as a loading control. Three independent experiments. **B.** Correlation between CYCLON protein levels (quantified from panel A) and dependency score (derived from DepMap database). R^2^ was derived from a simple linear regression. **C.** Immunofluorescence analysis of endogenous CYCLON localization in the pan-cancer cell line panel. Scale bar = 10 µm; DLBCL cells by confocal microscopy, scale bar = 5 µm. Two independent experiments.

**Supplementary Figure 2:**
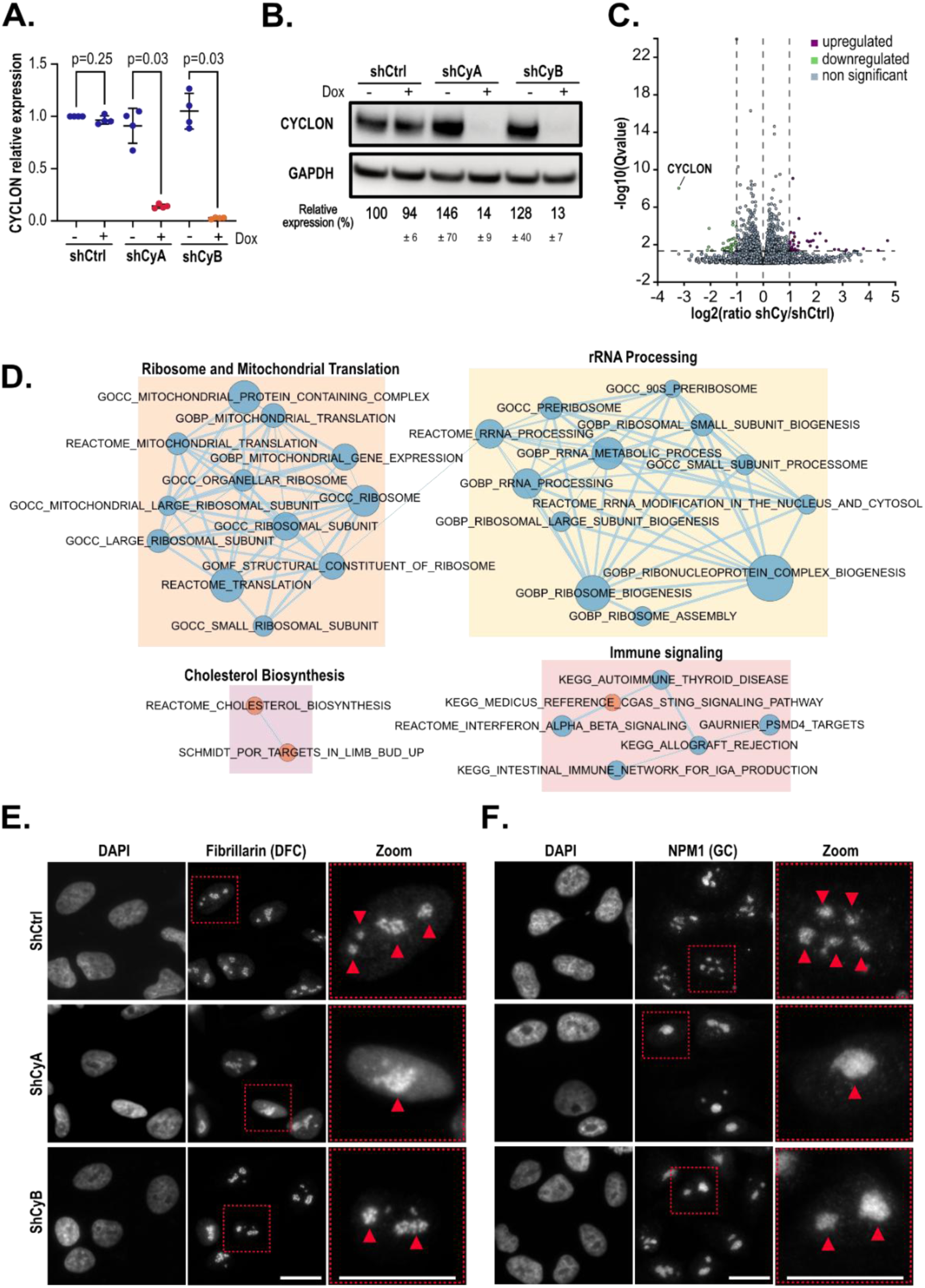
Validation and characterization of CYCLON doxycycline-inducible shRNA knock-down in HeLa cells. **A.** RT-qPCR analysis of CYCLON mRNA levels upon CYCLON silencing after 96h doxycycline induction. GAPDH and HSPC3 gene expression levels were used for normalization. Data are presented as individual values and mean ± SD. p values were derived from a Mann-Whitney Test) (n = 4 independent inductions). **B.** Western blot evaluation of CYCLON protein levels upon CYCLON silencing in cells transduced with a control (shCtrl) and CYCLON targeting shRNA (shCyA and shCyB) after 96h of doxycycline induction. GAPDH levels were used for normalization (n = 3 independent inductions). **C.** Volcano plot showing the differentially expressed genes between CYCLON KD (shCyA and shCyB) compared to shCtrl using the DeSeq2 pipeline **D**. Enrichment map of significantly enriched pathways identified by GSEA analysis on transcriptomic data (showed in panel C). Nodes represent gene sets, with red indicating positive normalized enrichment scores (NES) in the shCtrl condition and blue indicating positive NES in the shCy condition. **E-F.** Representative immunofluorescence images of Fibrillarin (E) and NPM1 (F) in cells transduced with a control (shCtrl) and CYCLON targeting shRNA (shCyA and shCyB) and DAPI nuclear staining, scale bar 20 µm, n = 30 images with 5-10 cells each, three independent experiments. The red arrows indicate morphological differences between shCy and shCtrl cells, while the dotted squares highlight the zoomed-in regions.

**Supplementary Figure 3.**
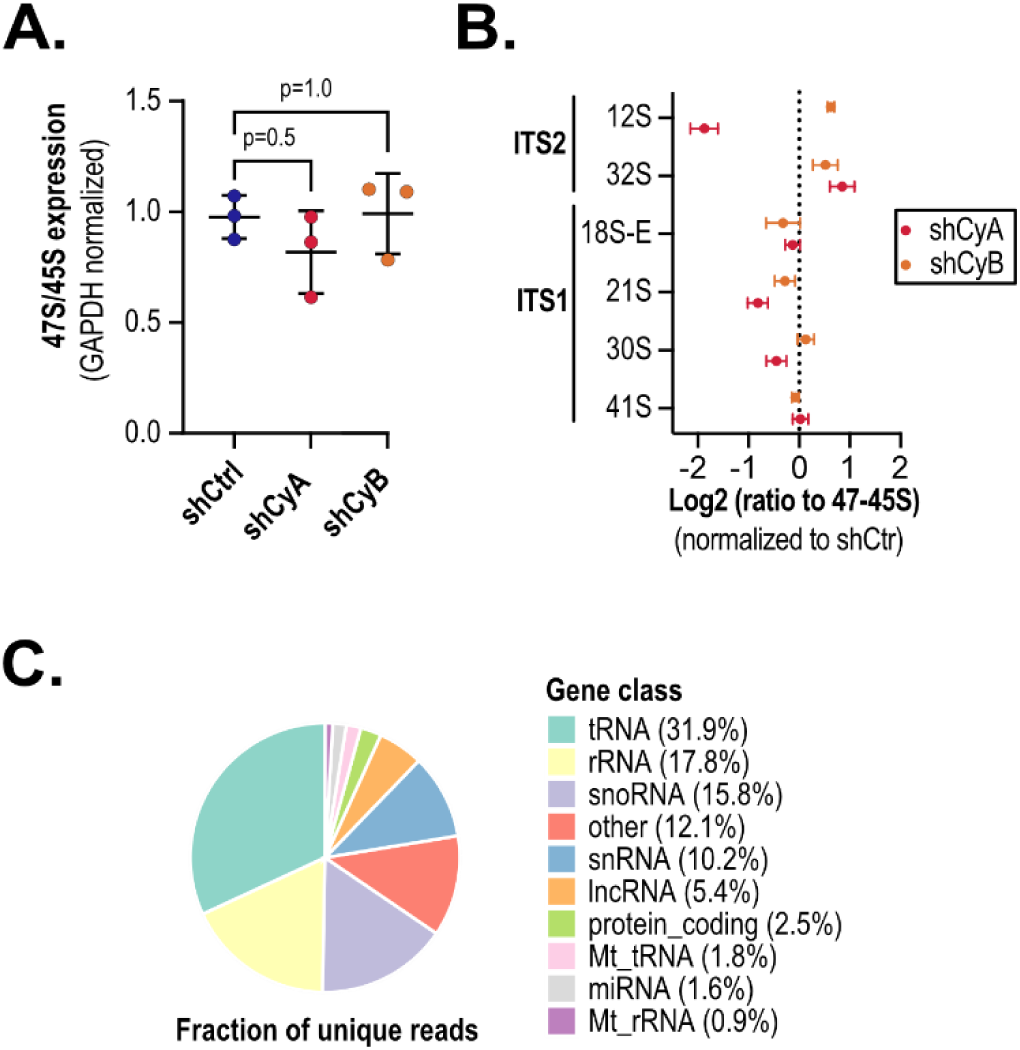
CYCLON plays a role in ribosome biogenesis in HeLa cells. **A.** RT-qPCR analysis of 45S precursor rRNA expression normalized to GAPDH in cells transduced with a control (shCtrl) and CYCLON targeting shRNA (shCyA et shCyB) after a 96h doxycycline induction, n = 3, p values are derived from a Kruskal-Wallis test followed by uncorrected Dunn’s test. **B.** Quantification of ratio between rRNA intermediates derived from Northern blot analyses in Fig 3B as indicated, n = 3, p values are derived from a Kruskal-Wallis test followed by uncorrected Dunn’s test. **C.** Fraction of unique RNA mapped species identified interacting with CYCLON in fCRAC analysis.

**Supplementary Figure 4.**
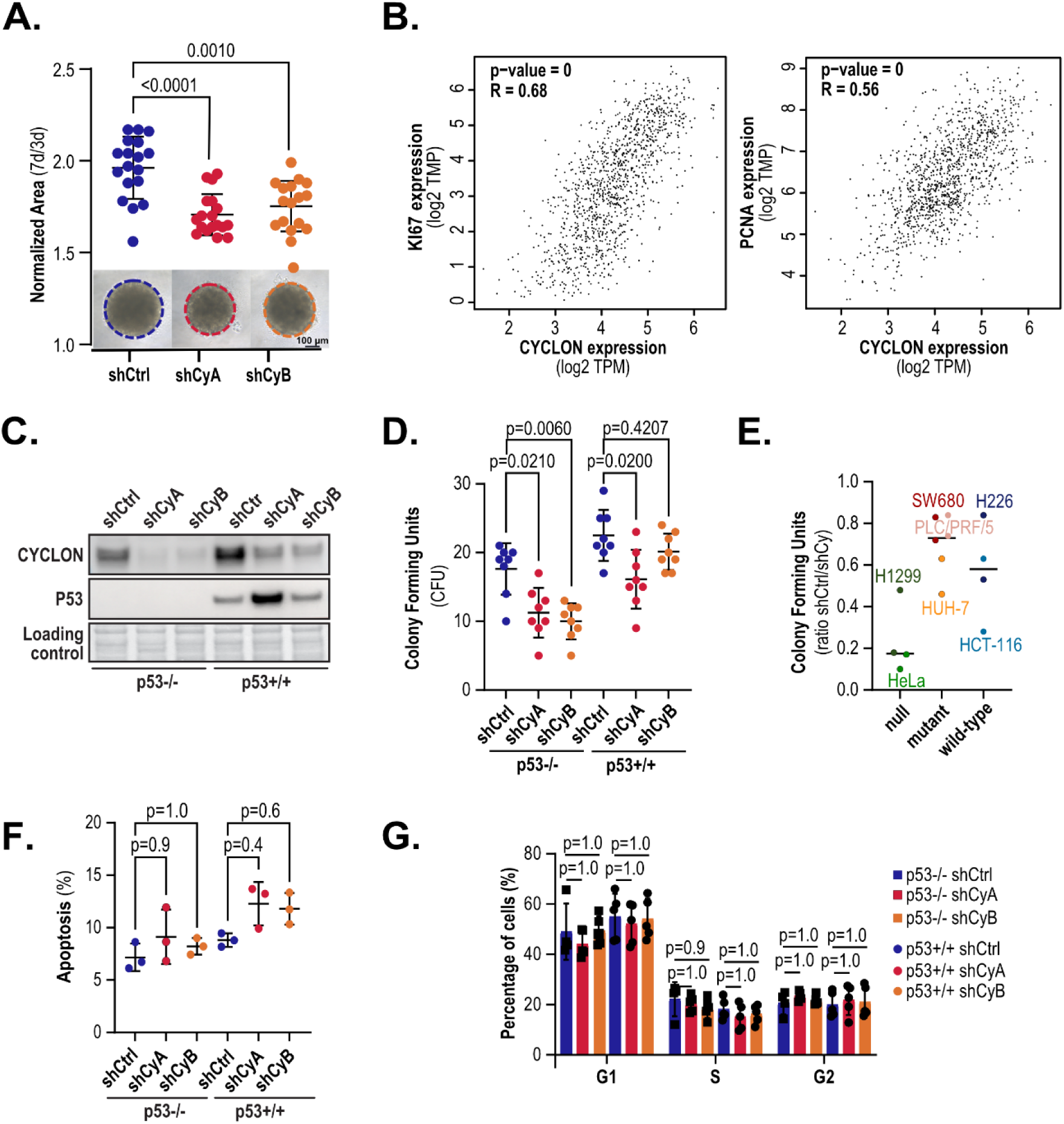
CYCLON regulates cell proliferation and clonogenic potential in pan-cancer manner independently of TP53 status. **A.** (up) Spheroid area 7 days after seeding of HeLa cells transduced with a control (shCtrl) and CYCLON targeting shRNA (shCyA et shCyB), day 7 area was normalized to day 3 area. n = 18, three independent experiments, (bottom) representative images at day 7, scale bar = 100 µm. p values were derived from a Brown-Forsythe test followed by uncorrected Dunnett’s test. **B.** Correlation plot between CYLON, Ki-67 and PCNA gene expression in TCGA COAD, DLBC, LIHC and LUSC databases. Plots were generated sing GEPIA2 interface (Pearson correlation test). **C.** Western blot analysis of p53 and CYCLON in HCT116 p53−/− and HCT116 p53+/+ transduced with a control (shCtrl) and CYCLON targeting shRNA (shCyA et shCyB). Ponceau staining was used as a loading control. **D.** Clonogenic assay in HCT116 p53−/− and HCT116 p53+/+ transduced with a control (shCtrl) and CYCLON targeting shRNA (shCyA et shCyB). **E.** Decrease in clonogenic potential in cell line models with null, mutant or wild-type p53 background as indicates. **F.** Apoptosis rate in HCT116 p53−/− and HCT116 p53+/+ cells transduced with a control (shCtrl) and CYCLON targeting shRNA (shCyA et shCyB), n = 6, six independent experiments. **G.** Cell cycle analysis performed in HeLa cells transduced with a control (shCtrl) and CYCLON targeting shRNA (shCyA et shCyB), n = 3, three independent experiments. P values are derived from Brown-Forsythe test followed by uncorrected Dunnett’s test.

## Bibliography

1. Lenz, L.S., Faccioni, J.L., Bracco, P.A., Santos, J.A.F., Pereira, L.C., Buss, J.H., Tamborindeguy, M.T., Torgo, D., Monteiro, T., Mantovani, G.B., et al. (2021) Cancer Cell Fitness Is Dynamic. Cancer Research, 81, 1040–1051.

2. Somarelli, J.A. (2021) The Hallmarks of Cancer as Ecologically Driven Phenotypes. Front. Ecol. Evol., 9, 661583.

3. Rozhok, A.I. and DeGregori, J. (2015) Toward an evolutionary model of cancer: Considering the mechanisms that govern the fate of somatic mutations. Proc. Natl. Acad. Sci. U.S.A., 112, 8914–8921.

4. Saint Fleur, S., Hoshino, A., Kondo, K., Egawa, T. and Fujii, H. (2009) Regulation of Fas-mediated immune homeostasis by an activation-induced protein, Cyclon. Blood, 114, 1355–1365.

5. Stamatiou, K., Chmielewska, A., Ohta, S., Earnshaw, W.C. and Vagnarelli, P. (2023) CCDC86 is a novel Ki-67-interacting protein important for cell division. Journal of Cell Science, 136, jcs260391.

6. Tsherniak, A., Vazquez, F., Montgomery, P.G., Weir, B.A., Kryukov, G., Cowley, G.S., Gill, S., Harrington, W.F., Pantel, S., Krill-Burger, J.M., et al. (2017) Defining a Cancer Dependency Map. Cell, 170, 564–576.e16.

7. Emadali, A., Rousseaux, S., Bruder-Costa, J., Rome, C., Duley, S., Hamaidia, S., Betton, P., Debernardi, A., Leroux, D., Bernay, B., et al. (2013) Identification of a novel BET bromodomain inhibitor-sensitive, gene regulatory circuit that controls Rituximab response and tumour growth in aggressive lymphoid cancers. EMBO Mol Med, 5, 1180–1195.

8. Bouroumeau, A., Bussot, L., Sartelet, H., Fournier, C., Betton-Fraisse, P., Col, E., David-Boudet, L., McLeer, A., Lefebvre, C., Raskovalova, T., et al. (2021) Extranucleolar CYCLON Staining Pattern Is Strongly Associated to Relapse/Refractory Disease in R-CHOP–treated DLBCL. Hemasphere, 5, e598.

9. Wang, Z., Zhou, T., Chen, X., Zhu, X., Liao, B., Liu, J., Li, S., Tan, T. and Liu, Y. (2024) CCDC86 promotes the aggressive behavior of nasopharyngeal carcinoma by positively regulating EGFR and activating the PI3K/Akt signaling. neo, 70, 761–776.

10. Hoshino, A. and Fujii, H. (2007) Redundant promoter elements mediate IL-3-induced expression of a novel cytokine-inducible gene, *cyclon*. FEBS Letters, 581, 975–980.

11. Dominguez-Sola, D., Victora, G.D., Ying, C.Y., Phan, R.T., Saito, M., Nussenzweig, M.C. and Dalla-Favera, R. (2012) The proto-oncogene MYC is required for selection in the germinal center and cyclic reentry. Nat Immunol, 13, 1083–1091.

12. Moy, T.I., Boettner, D., Rhodes, J.C., Silver, P.A. and Askew, D.S. (2002) Identification of a role for Saccharomyces cerevisiae Cgr1p in pre-rRNA processing and 60S ribosome subunit synthesis. Microbiology, 148, 1081–1090.

13. Bouroumeau, A., Bussot, L., Hamaidia, S., Garcìa-Sandoval, A., Bergan-Dahl, A., Betton-Fraisse, P., Duley, S., Fournier, C., Aucagne, R., Adrait, A., et al. (2021) CYCLON and NPM1 Cooperate within an Oncogenic Network Predictive of R-CHOP Response in DLBCL. Cancers (Basel), 13, 5900.

14. Baltz, A.G., Munschauer, M., Schwanhäusser, B., Vasile, A., Murakawa, Y., Schueler, M., Youngs, N., Penfold-Brown, D., Drew, K., Milek, M., et al. (2012) The mRNA-Bound Proteome and Its Global Occupancy Profile on Protein-Coding Transcripts. Molecular Cell, 46, 674–690.

15. Becker, J.S., McCarthy, R.L., Sidoli, S., Donahue, G., Kaeding, K.E., He, Z., Lin, S., Garcia, B.A. and Zaret, K.S. (2017) Genomic and Proteomic Resolution of Heterochromatin and Its Restriction of Alternate Fate Genes. Molecular Cell, 68, 1023–1037.e15.

16. Beckmann, B.M., Horos, R., Fischer, B., Castello, A., Eichelbaum, K., Alleaume, A.-M., Schwarzl, T., Curk, T., Foehr, S., Huber, W., et al. (2015) The RNA-binding proteomes from yeast to man harbour conserved enigmRBPs. Nat Commun, 6, 10127.

17. Castello, A., Fischer, B., Eichelbaum, K., Horos, R., Beckmann, B.M., Strein, C., Davey, N.E., Humphreys, D.T., Preiss, T., Steinmetz, L.M., et al. (2012) Insights into RNA Biology from an Atlas of Mammalian mRNA-Binding Proteins. Cell, 149, 1393–1406.

18. Dürnberger, G., Bürckstümmer, T., Huber, K., Giambruno, R., Doerks, T., Karayel, E., Burkard, T.R., Kaupe, I., Müller, A.C., Schönegger, A., et al. (2013) Experimental characterization of the human non-sequence-specific nucleic acid interactome. Genome Biol, 14, R81.

19. Ginno, P.A., Burger, L., Seebacher, J., Iesmantavicius, V. and Schübeler, D. (2018) Cell cycle-resolved chromatin proteomics reveals the extent of mitotic preservation of the genomic regulatory landscape. Nat Commun, 9, 4048.

20. Queiroz, R.M.L., Smith, T., Villanueva, E., Marti-Solano, M., Monti, M., Pizzinga, M., Mirea, D.-M., Ramakrishna, M., Harvey, R.F., Dezi, V., et al. (2019) Comprehensive identification of RNA–protein interactions in any organism using orthogonal organic phase separation (OOPS). Nat Biotechnol, 37, 169–178.

21. Trendel, J., Schwarzl, T., Horos, R., Prakash, A., Bateman, A., Hentze, M.W. and Krijgsveld, J. (2019) The Human RNA-Binding Proteome and Its Dynamics during Translational Arrest. Cell, 176, 391–403.e19.

22. Boisvert, F.-M., van Koningsbruggen, S., Navascués, J. and Lamond, A.I. (2007) The multifunctional nucleolus. Nat Rev Mol Cell Biol, 8, 574–585.

23. Frottin, F., Schueder, F., Tiwary, S., Gupta, R., Körner, R., Schlichthaerle, T., Cox, J., Jungmann, R., Hartl, F.U. and Hipp, M.S. (2019) The nucleolus functions as a phase-separated protein quality control compartment. Science, 365, 342–347.

24. Cuylen, S., Blaukopf, C., Politi, A.Z., Müller-Reichert, T., Neumann, B., Poser, I., Ellenberg, J., Hyman, A.A. and Gerlich, D.W. (2016) Ki-67 acts as a biological surfactant to disperse mitotic chromosomes. Nature, 535, 308–312.

25. Sobecki, M., Mrouj, K., Camasses, A., Parisis, N., Nicolas, E., Llères, D., Gerbe, F., Prieto, S., Krasinska, L., David, A., et al. (2016) The cell proliferation antigen Ki-67 organises heterochromatin. eLife, 5, e13722.

26. Orsolic, I., Jurada, D., Pullen, N., Oren, M., Eliopoulos, A.G. and Volarevic, S. (2016) The relationship between the nucleolus and cancer: Current evidence and emerging paradigms. Seminars in Cancer Biology, 37-38, 36–50.

27. Wiederschain, D., Susan, W., Chen, L., Loo, A., Yang, G., Huang, A., Chen, Y., Caponigro, G., Yao, Y., Lengauer, C., et al. (2009) Single-vector inducible lentiviral RNAi system for oncology target validation. Cell Cycle, 8, 498–504.

28. Dobin, A., Davis, C.A., Schlesinger, F., Drenkow, J., Zaleski, C., Jha, S., Batut, P., Chaisson, M. and Gingeras, T.R. (2013) STAR: ultrafast universal RNA-seq aligner. Bioinformatics, 29, 15–21.

29. Putri, G.H., Anders, S., Pyl, P.T., Pimanda, J.E. and Zanini, F. (2022) Analysing high-throughput sequencing data in Python with HTSeq 2.0. Bioinformatics, 38, 2943–2945.

30. Varet, H., Brillet-Guéguen, L., Coppée, J.-Y. and Dillies, M.-A. (2016) SARTools: A DESeq2– and EdgeR-Based R Pipeline for Comprehensive Differential Analysis of RNA-Seq Data. PLOS ONE, 11, e0157022.

31. Love, M.I., Huber, W. and Anders, S. (2014) Moderated estimation of fold change and dispersion for RNA-seq data with DESeq2. Genome Biol, 15, 1–21.

32. Subramanian, A., Tamayo, P., Mootha, V.K., Mukherjee, S., Ebert, B.L., Gillette, M.A., Paulovich, A., Pomeroy, S.L., Golub, T.R., Lander, E.S., et al. (2005) Gene set enrichment analysis: A knowledge-based approach for interpreting genome-wide expression profiles. Proc Natl Acad Sci U S A, 102, 15545–15550.

33. Paraqindes, H., Mourksi, N.-E.-H., Ballesta, S., Hedjam, J., Bourdelais, F., Fenouil, T., Picart, T., Catez, F., Combe, T., Ferrari, A., et al. (2023) *Isocitrate dehydrogenase* wt and IDHmut adult-type diffuse gliomas display distinct alterations in ribosome biogenesis and 2’O-methylation of ribosomal RNA. Neuro-Oncology, 25, 2191–2206.

34. Stenström, L., Mahdessian, D., Gnann, C., Cesnik, A.J., Ouyang, W., Leonetti, M.D., Uhlén, M., Cuylen-Haering, S., Thul, P.J. and Lundberg, E. (2020) Mapping the nucleolar proteome reveals a spatiotemporal organization related to intrinsic protein disorder. Molecular Systems Biology, 16, e9469.

35. Ugrinova, I., Monier, K., Ivaldi, C., Thiry, M., Storck, S., Mongelard, F. and Bouvet, P. (2007) Inactivation of nucleolin leads to nucleolar disruption, cell cycle arrest and defects in centrosome duplication. BMC Mol Biol, 8, 66.

36. Huttlin, E.L., Ting, L., Bruckner, R.J., Gebreab, F., Gygi, M.P., Szpyt, J., Tam, S., Zarraga, G., Colby, G., Baltier, K., et al. (2015) The BioPlex Network: A Systematic Exploration of the Human Interactome. Cell, 162, 425–440.

37. Liu, X., Salokas, K., Tamene, F., Jiu, Y., Weldatsadik, R.G., Öhman, T. and Varjosalo, M. (2018) An AP-MS– and BioID-compatible MAC-tag enables comprehensive mapping of protein interactions and subcellular localizations. Nat Commun, 9, 1188.

38. Cho, N.H., Cheveralls, K.C., Brunner, A.-D., Kim, K., Michaelis, A.C., Raghavan, P., Kobayashi, H., Savy, L., Li, J.Y., Canaj, H., et al. (2022) OpenCell: Endogenous tagging for the cartography of human cellular organization. Science, 375, eabi6983.

39. Hein, M.Y., Hubner, N.C., Poser, I., Cox, J., Nagaraj, N., Toyoda, Y., Gak, I.A., Weisswange, I., Mansfeld, J., Buchholz, F., et al. (2015) A Human Interactome in Three Quantitative Dimensions Organized by Stoichiometries and Abundances. Cell, 163, 712–723.

40. Daniely, Y., Dimitrova, D.D. and Borowiec, J.A. (2002) Stress-Dependent Nucleolin Mobilization Mediated by p53-Nucleolin Complex Formation. Molecular and Cellular Biology, 22, 6014.

41. Hirai, Y., Louvet, E., Oda, T., Kumeta, M., Watanabe, Y., Horigome, T. and Takeyasu, K. (2013) Nucleolar scaffold protein, WDR 46, determines the granular compartmental localization of nucleolin and DDX 21. Genes to Cells, 18, 780–797.

42. Potapova, T.A., Unruh, J.R., Conkright-Fincham, J., Banks, C.A., Florens, L., Schneider, D.A. and Gerton, J.L. (2023) Distinct states of nucleolar stress induced by anticancer drugs. eLife, 12, RP88799.

43. Koh, C.M., Gurel, B., Sutcliffe, S., Aryee, M.J., Schultz, D., Iwata, T., Uemura, M., Zeller, K.I., Anele, U., Zheng, Q., et al. (2011) Alterations in Nucleolar Structure and Gene Expression Programs in Prostatic Neoplasia Are Driven by the MYC Oncogene. The American Journal of Pathology, 178, 1824–1834.

44. Robertson, N., Shchepachev, V., Wright, D., Turowski, T.W., Spanos, C., Helwak, A., Zamoyska, R. and Tollervey, D. (2022) A disease-linked lncRNA mutation in RNase MRP inhibits ribosome synthesis. Nat Commun, 13, 649.

45. Granneman, S., Petfalski, E. and Tollervey, D. (2011) A cluster of ribosome synthesis factors regulate pre-rRNA folding and 5.8S rRNA maturation by the Rat1 exonuclease: Pre-60S ribosome synthesis. The EMBO Journal, 30, 4006–4019.

46. Lebaron, S., Segerstolpe, Å., French, S.L., Dudnakova, T., de lima Alves, F., Granneman, S., Rappsilber, J., Beyer, A.L., Wieslander, L. and Tollervey, D. (2013) Rrp5 Binding at Multiple Sites Coordinates Pre-rRNA Processing and Assembly. Molecular Cell, 52, 707–719.

47. Turowski, T.W., Petfalski, E., Goddard, B.D., French, S.L., Helwak, A. and Tollervey, D. (2020) Nascent Transcript Folding Plays a Major Role in Determining RNA Polymerase Elongation Rates. Molecular Cell, 79, 488–503.e11.

48. Goldfarb, K.C. and Cech, T.R. (2017) Targeted CRISPR disruption reveals a role for RNase MRP RNA in human preribosomal RNA processing. Genes Dev., 31, 59–71.

49. Hannan, K.M., Soo, P., Wong, M.S., Lee, J.K., Hein, N., Poh, P., Wysoke, K.D., Williams, T.D., Montellese, C., Smith, L.K., et al. (2022) Nuclear stabilization of p53 requires a functional nucleolar surveillance pathway. Cell Reports, 41, 111571.

50. James, A., Wang, Y., Raje, H., Rosby, R. and DiMario, P. (2014) Nucleolar stress with and without p53. Nucleus, 5, 402–426.

51. Krüger, T. and Scheer, U. (2010) p53 localizes to intranucleolar regions distinct from the ribosome production compartments. Journal of Cell Science, 123, 1203–1208.

52. Jordan, E.G. and Mcgovern, J.H. (1981) The quantitative relationship of the fibrillar centres and other nucleolar components to changes in growth conditions, serum deprivation and low doses of actinomycin D in cultured diploid human fibroblasts (strain mrc-5). Journal of Cell Science, 52, 373–389.

53. Derenzini, M., Montanaro, L. and Treré, D. (2009) What the nucleolus says to a tumour pathologist. Histopathology, 54, 753–762.

54. Lafontaine, D.L.J., Riback, J.A., Bascetin, R. and Brangwynne, C.P. (2021) The nucleolus as a multiphase liquid condensate. Nat Rev Mol Cell Biol, 22, 165–182.

55. Nicolas, E., Parisot, P., Pinto-Monteiro, C., De Walque, R., De Vleeschouwer, C. and Lafontaine, D.L.J. (2016) Involvement of human ribosomal proteins in nucleolar structure and p53-dependent nucleolar stress. Nat Commun, 7, 11390.

56. Stamatopoulou, V., Parisot, P., De Vleeschouwer, C. and Lafontaine, D.L.J. (2018) Use of the iNo score to discriminate normal from altered nucleolar morphology, with applications in basic cell biology and potential in human disease diagnostics. Nature Protocols, 13, 2387–2406.

57. Amin, M.A., Matsunaga, S., Uchiyama, S. and Fukui, K. (2008) Depletion of nucleophosmin leads to distortion of nucleolar and nuclear structures in HeLa cells. Biochemical Journal, 415, 345–351.

58. Erales, J., Marchand, V., Panthu, B., Gillot, S., Belin, S., Ghayad, S.E., Garcia, M., Laforêts, F., Marcel, V., Baudin-Baillieu, A., et al. (2017) Evidence for rRNA 2′-O-methylation plasticity: Control of intrinsic translational capabilities of human ribosomes. Proc. Natl. Acad. Sci. U.S.A., 114, 12934–12939.

59. Replogle, J.M., Saunders, R.A., Pogson, A.N., Hussmann, J.A., Lenail, A., Guna, A., Mascibroda, L., Wagner, E.J., Adelman, K., Lithwick-Yanai, G., et al. (2022) Mapping information-rich genotype-phenotype landscapes with genome-scale Perturb-seq. Cell, 185, 2559–2575.e28.

60. Vanden Broeck, A. and Klinge, S. (2023) Principles of human pre-60 S biogenesis. Science, 381, eadh3892.

61. Zhang, Y., Liang, X., Luo, S., Chen, Y., Li, Y., Ma, C., Li, N. and Gao, N. (2023) Visualizing the nucleoplasmic maturation of human pre-60S ribosomal particles. Cell Res, 33, 867–878.

62. Holmberg Olausson, K., Nistér, M. and Lindström, M.S. (2012) p53 –Dependent and –Independent Nucleolar Stress Responses. Cells, 1, 774–798.

63. Moraleva, А., Antipova, N., Pavlov, P., Dobrochaeva, K. and Rubtsov, Y. Nucleolus as a cornerstone linking proliferation and metabolism to cellular responses to stress: involvement of transcription factors MYC and p53. Front Mol Biosci, 13, 1749992.

64. Chen, D., Lu, S., Huang, K., Pearson, J.D., Pacal, M., Peidis, P., McCurdy, S., Yu, T., Sangwan, M., Nguyen, A., et al. (2025) Cell cycle duration determines oncogenic transformation capacity. Nature, 641, 1309–1318.

65. Mrouj, K., Andrés-Sánchez, N., Dubra, G., Singh, P., Sobecki, M., Chahar, D., Al Ghoul, E., Aznar, A.B., Prieto, S., Pirot, N., et al. (2021) Ki-67 regulates global gene expression and promotes sequential stages of carcinogenesis. Proc. Natl. Acad. Sci. U.S.A., 118, e2026507118.

